# Colorectal ALOX15 as a host factor determinant of EPA and DHA effects on colorectal carcinogenesis

**DOI:** 10.1101/2024.05.02.592224

**Authors:** Xiangsheng Zuo, Yoshiyuki Kiyasu, Yi Liu, Yasunori Deguchi, Fuyao Liu, Micheline Moussalli, Lin Tan, Bo Wei, Daoyan Wei, Peiying Yang, Imad Shureiqi

**Author notes:** Correspondence (I.S.), Division of Hematology and Oncology, Department of Internal Medicine, and Rogel Cancer Center, University of Michigan, Ann Arbor, MI48109. These authors contributed equally to this manuscript (Xiangsheng Zuo, Yoshiyuki Kiyasu, Yi Liu). **Conflicts of interest:** The authors declare no conflicts.

## Abstract

Eicosapentaenoic acid (EPA) and docosahexaenoic acid (DHA), omega-3 polyunsaturated fatty acids (ω-3 PUFAs) derived from fish oil, are widely used as dietary supplements and FDA-approved treatments for hypertriglyceridemia. However, studies investigating the effects of EPA and DHA on colorectal carcinogenesis (CRC) have yielded conflicting results. The factors that determine these discrepant results remain unknown. Resolvins, oxidative metabolites of EPA and DHA, inhibit key pro-tumorigenic cytokine and chemokine signaling of colorectal cancer (e.g., IL-6, IL-1β, and CCL2). 15-lipoxygenase-1 (ALOX15), a critical enzyme for resolvin generation is commonly lost during human CRC. Whether ALOX15 expression, as a host factor, modulates the effects of EPA and DHA on CRC remains unknown. Therefore, we evaluated the effects of ALOX15 transgenic expression in colonic epithelial cells on resolvin generation by EPA and DHA and CRC in mouse models representative of human CRC. Our results revealed that 1) EPA and DHA effects on CRC were diverse, ranging from suppressive to promotive, and these effects were occasionally altered by the formulations of EPA and DHA (free fatty acid, ethyl ester, triglyceride); 2) EPA and DHA uniformly suppressed CRC in the presence of intestinal ALOX15 transgenic expression, which induced the production of resolvins, decreased colonic CCL3-5 and CXCL-5 expression and tumor associated macrophages while increasing CD8 T cell abundance in tumor microenvironment; and 3) RvD5, the predominant resolvin produced by ALOX15, inhibited macrophage generation of pro-tumorigenic cytokines. These findings demonstrate the significance of intestinal ALOX15 expression as a host factor in determining the effects of EPA and DHA on CRC.

**Significance:** Eicosapentaenoic acid (EPA) and docosahexaenoic acid (DHA) are widely used as dietary supplements and FDA-approved treatments for hypertriglyceridemia. Studies of EPA and DHA effects on colorectal carcinogenesis (CRC) have revealed inconsistencies; factors determining the direction of their impact on CRC have remained unidentified. Our data show that EPA and DHA effects on CRC were divergent and occasionally influenced by their formulations. More importantly, intestinal 15-lipoxgenase-1 (ALOX15) expression modulated EPA and DHA effects on CRC, leading to their consistent suppression of CRC. ALOX15 promoted EPA and DHA oxidative metabolism to generate resolvins, which inhibited key pro-tumorigenic inflammatory cytokines and chemokines, including IL-6. IL-1β, and CCL2. ALOX15 is therefore an important host factor in determining EPA and DHA effects on CRC.

## INTRODUCTION

Fish oil and its major omega-3 polyunsaturated fatty acid (ω-3 PUFAs) derivatives, eicosapentaenoic acid (EPA) and docosahexaenoic acid (DHA), are widely promoted as remedies to prevent major chronic diseases, including cancers. The commonly used formulations of EPA and DHA are free fatty acids (FFA), ethyl esters (EE), and triglycerides (TG). According to the 2012 National Health Interview Survey, approximately 19 million American adults consume dietary fish oil supplements (1). Additionally, Lovaza, a combination of EPA-EE and DHA-EE, and Vascepa, a highly purified EPA-EE formulation, are approved by the Food and Drug Administration (FDA) for hypertriglyceridemia treatment.

Western diets commonly have low levels of EPA and DHA (2, 3). Studies investigating the link between fish oil, EPA, DHA, and cancers, including colorectal carcinogenesis (CRC), have largely focused on the effects of EPA and DHA supplementation. However, the effects of EPA and DHA supplementation on cancer risk have been inconsistent (4, 5). For example, while fish oil was reported to suppress CRC in various preclinical rodent studies (6–8), it was also reported to promote CRC, especially with DHA– enriched formulations (9, 10). EPA inhibited CRC in preclinical models (11, 12) and in a randomized clinical trial of familial adenomatous polyposis patients (13), but failed to reduce sporadic polyp recurrence in a subsequent larger randomized clinical study (14). Other clinical studies have also shown variable effects, including beneficial, null, and harmful effects, of EPA and DHA supplements on cancer risk (13–18). These inconsistent results raise the question of whether the mere availability of EPA and DHA is sufficient to reproducibly suppress CRC.

Host enzymatic oxidation of EPA and DHA produces biologically active oxidative metabolites (e.g., resolvins) that can potentially affect carcinogenesis. Resolvins have been reported to repress the production of pro-inflammatory cytokines and chemokines (e.g., IL-6, IL-1β, CCL2) that promote CRC (19–21). 15-lipoxygenase-1 (ALOX15) catalyzes the oxidative metabolism of DHA to 17-S-HpDHA, a precursor of the resolvin D series (e.g., RvD1-5) (22, 23). 15-LOX-like function of aspirin-acetylated cyclooxygenase-2 (COX-2) enzyme catalyzes the oxidative metabolism of EPA to 18-HEPE, a precursor of the resolvin E series (e.g., RvE1-3) (24). 5-lipoxygenase (5-LOX) promotes subsequent metabolism of resolvin precursors to generate resolvins (25). Whereas 5-LOX is upregulated during carcinogenesis (26), ALOX15 is downregulated in various major human cancers, especially in colorectal cancer (27–29) where it is lost in up to 67% of high-grade colonic adenomas and 100% of human invasive colorectal cancers (27, 30). Nevertheless, published data regarding the mechanistic significance of ALOX15 to EPA and DHA effects on carcinogenesis are limited to a report of in vitro experiments showing that ALOX15 is required for DHA induction of apoptosis in prostate cancer cells (31). Whether ALOX15 loss during CRC affects resolvin generation from EPA and DHA and subsequently CRC remains largely unknown. The mouse homolog of human ALOX15 is 12/15 LOX, which is a hybrid enzyme that produces both of the ALOX15 and 12-S-LOX products (i.e. 13-S-HODE and 12-S-HETE) that have opposing effects on carcinogenesis (32, 33). We therefore conducted the current studies to address this knowledge gap by examining the effects of transgenic expression of human ALOX15 in mouse intestines on resolvin generation from the three common EPA and DHA formulations, as well as the subsequent impact on CRC, using multiple in vivo experimental mouse CRC models that are representative of human CRC.

## RESULTS

### ALOX15 modulates the effects of dietary fish oil, EPA-FFA, and DHA-FFA on CRC in multiple CRC mouse models

To investigate the impact of ALOX15 on oxidative metabolism of EPA and DHA and subsequent CRC, we first examined the effects of intestinal ALOX15 transgenic expression in ALOX15-Gut mice fed a fish oil supplemented diet on AOM/DSS–induced colitis-associated colorectal cancer (CAC). ALOX15-Gut and wild-type (WT) littermates were treated with azoxymethane (AOM) and dextran sodium sulfate (DSS) while they were fed a fish oil diet containing 11.5% menhaden oil or an isocaloric control diet (34) (Supplementary Table 1). Unexpectedly, WT mice fed with the Menhaden fish oil diet developed higher colonic tumor multiplicity than the same mice fed control diet (Fig.1A and B, Supplementary Table 2). Intestinal ALOX15 transgenic expression significantly decreased colonic tumor multiplicity in ALOX15-Gut mice fed either fish oil or control diet when compared to the same diet–fed WT littermates (Fig. 1A and B, Supplementary Table 2).

**Figure 1.**
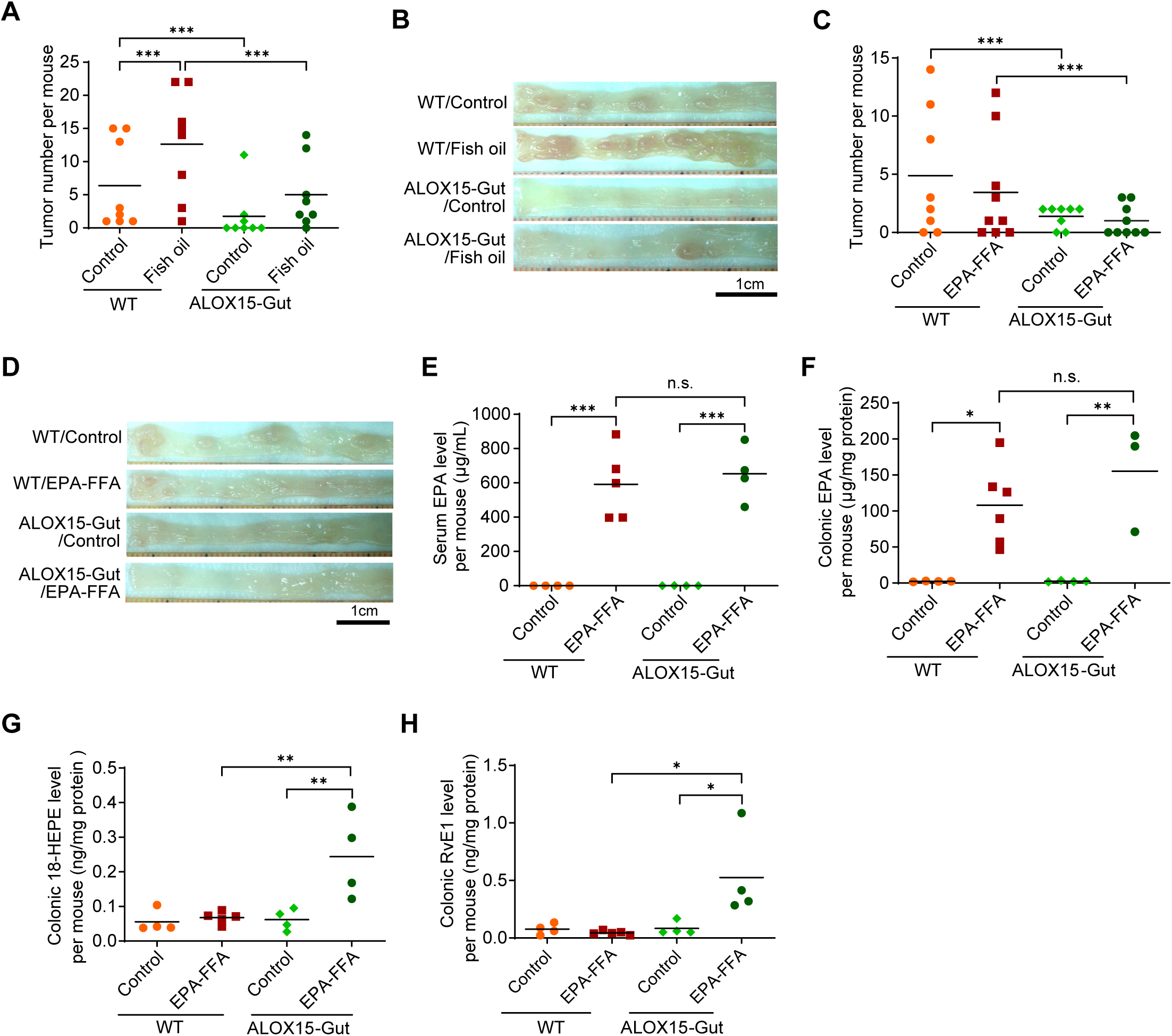
ALOX15 modulates the effects of fish oil and EPA-FFA on AOM/DSS–induced CAC and the production of resolvins. (A, B) ALOX15-Gut mice and their WT littermates fed 11.5% menhaden fish oil or control diet were treated with AOM and DSS to induce CAC as described in the Methods section (n = 8 mice per group). Total colonic tumor number per mouse (A) and the representative gross colon images (B) for the indicated mouse groups are shown. (C-H) ALOX15-Gut mice and their WT littermates fed 1% EPA-FFA or control diet were treated with AOM and DSS to induce CAC (n = 8-9 mice per group). (C, D) Total colonic tumor number per mouse (C) and the representative gross colon images (D) for the indicated mouse groups are shown. (E-H) The levels of serum EPA (E), colonic EPA (F), colonic 18-HEPE (G), and colonic RvE1 (H) per mouse for the indicated mouse groups, measured by LC/MS/MS, are shown. Lines for panels A, C and E-H represent means. Poisson regression for panels A and C, and two-way ANOVA with Bonferroni adjustments for all multiple comparisons for panels E-H. *\* P < 0.05; ** P < 0.01; *** P <0.001;* and n.s.: non-significant.

We next evaluated the effects of ALOX15 and EPA-FFA on CAC using the same CAC mouse model. ALOX15-Gut and WT littermates were fed with 1% highly purified EPA-FFA or control diet (Supplementary Table 3). Colorectal tumor multiplicity trended to be lower in EPA-FFA diet–fed WT mice than in control diet–fed WT mice, although the difference failed to reach statistical significance (*P=* 0.14) (Fig. 1C and D). ALOX15-Gut mice fed the control diet or EPA-FFA diet had significantly lower tumor multiplicity than WT mice fed the same diet (Fig. 1C and D).

We then evaluated whether the transgenic expression of ALOX15 in the mice fed EPA-FFA diet modulated resolvin production as it suppressed CAC. EPA levels markedly increased in the sera and colonic epithelial cells in both WT and ALOX15-Gut mice when fed the EPA-FFA diet (Fig. 1E, F). Despite equal EPA bioavailability between ALOX15-Gut mice and WT control littermates, colonic 18-HEPE and RvE1 levels significantly increased in ALOX15-Gut mice fed EPA-FFA diet but not in the other mouse groups (Fig. 1G, H). These findings demonstrated that ALOX15 was necessary for resolvin generation from EPA-FFA. In follow-up independent experiments in an AOM–induced CRC mouse model, EPA-FFA or/and intestinal ALOX15 expression significantly reduced colorectal tumor volumes, and the tumor volumes in EPA-FFA diet–fed ALOX15-Gut mice trended to be lower than tumor volumes in EPA-FFA diet–fed WT (*P =* 0.1) and control diet–fed ALOX15-Gut mice (*P =* 0.07) (Supplementary Fig. 1).

To examine whether the ALOX15 effects are specific to EPA or CRC models induced by AOM /DSS or AOM, we evaluated the efffects of DHA-FFA on CRC using Apc^580mu^ mice with (Apc^580mu^-ALOX15-Gut) and without (Apc^580mu^) intestinal ALOX15 transgenic expression. DHA-FFA (1%) supplemented diet (Supplementary Table 3) significantly increased levels of serum DHA in Apc^580mu^ and Apc^580mu^-ALOX15-Gut mice (Supplementary Fig. 2A). However, colonic 17-HDHA and RvD2 levels increased in Apc^580mu^-ALOX15-Gut mice but not in Apc^580mu^ mice (Supplementary Fig. 2B, C). Dietary DHA-FFA significantly increased the multiplicity of large tumors (diameter > 4mm), but intestinal ALOX15 expression repressed this DHA’s effect (Supplementary Fig. 2D, E) and reduced the total tumor multiplicity in Apc^580mu^-ALOX15-Gut mice fed both control and DHA-FFA diet (Supplementary Fig. 2D, F). Taken together, the findings from the fish oil, EPA and DHA independent experiments indicated that fish oil and its derivatives had variable effects on CRC, and ALOX15 modulated their effects to uniformly suppress CRC.

### ALOX15 modulates the effects of dietary EPA-FFA, DHA-FFA and Lovaza on AOM–induced CRC

We next directly compared the effects of dietary EPA-FFA, DHA-FFA, and Lovaza on AOM–induced CRC in ALOX15-Gut mice and their WT littermates. Lovaza capsules contain EPA-EE (46.5%) and DHA-EE (37.5%), thus a 2.7% Lovaza diet contains dietary 1.3% EPA-EE and 1% DHA-EE. ALOX15-Gut mice and their WT littermates were treated with AOM (7.5 mg/kg) and fed 1% DHA-FFA, or 1% EPA-FFA, or 2.7% Lovaza or control diet (Supplementary Table 3). Overall, colorectal tumor multiplicity and volume trended to be lower in EPA-FFA diet– and Lovaza diet–fed mice than in control diet– or DHA-FFA diet–fed mice, especially in ALOX15-Gut mice (Fig. 2A, B, Supplementary Table 4). Lovaza diet–fed WT mice had significantly lower colorectal tumor numbers and volumes than control diet–fed WT mice; EPA-FFA–fed WT mice had similar difference trend, but it did not reach statistical significance. In contrast, colorectal tumor numbers and volumes trended to be higher in DHA-FFA diet–fed WT mice than in control diet–fed WT mice (Fig. 2A, B, Supplementary Tables 4). Intestinal ALOX15 transgenic expression reduced colonic tumor multiplicity and volumes across all diet conditions in ALOX15*-*Gut mice (Fig. 2A and B, Supplementary Table 4), and increased levels of colonic 18-HEPE, RvE1, and 17-HDHA in ALOX15*-*Gut mice when fed DHA-FFA, EPA-FFA, or Lovaza diet, and the levels of colonic RvD2 in ALOX15*-*Gut mice when fed control, DHA-FFA, or Lovaza diet (Fig. 2C-F).

**Figure 2.**
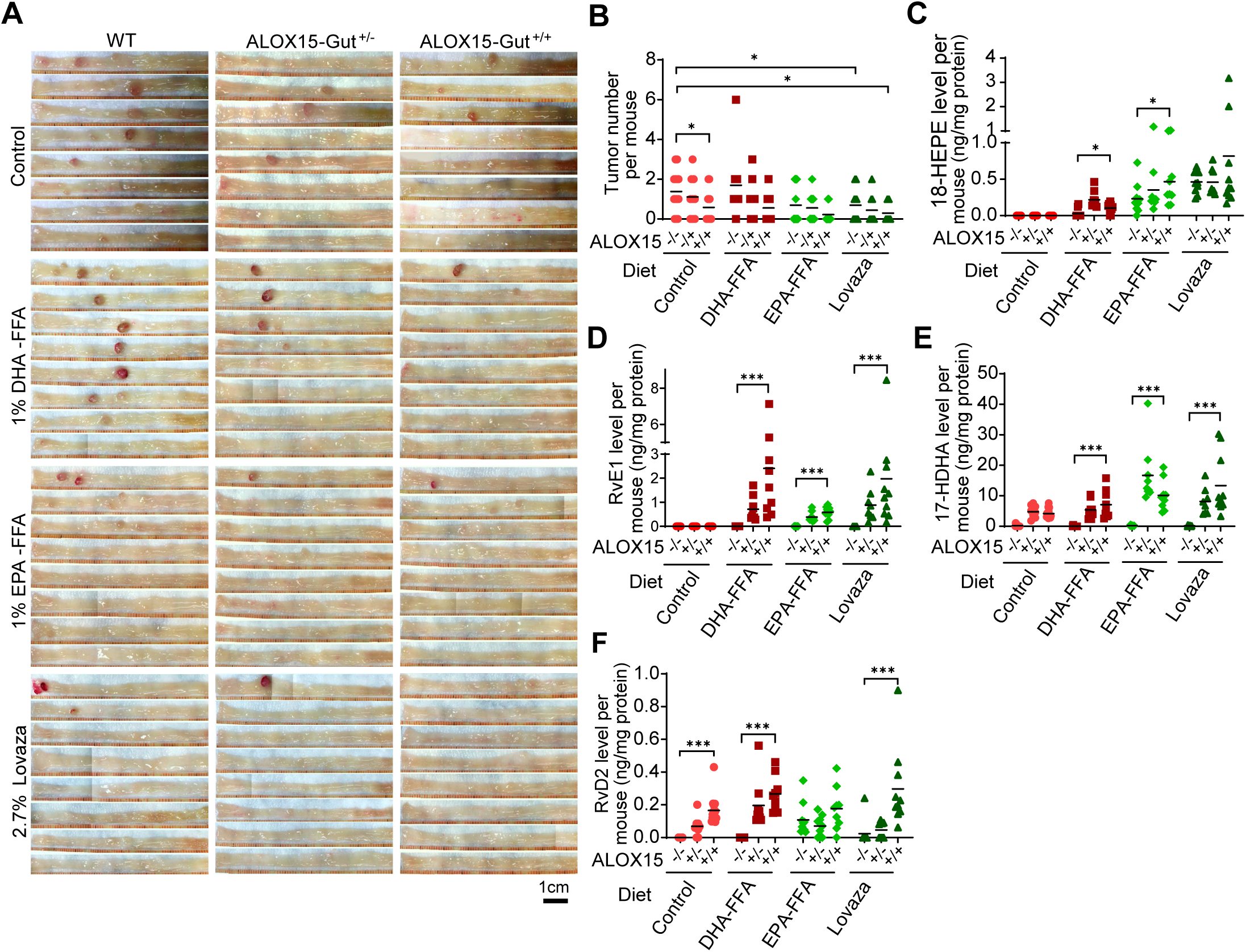
ALOX15 modulates the effects of dietary EPA-FFA, DHA-FFA and Lovaza on AOM– induced CRC and the production of resolvins. ALOX15-Gut mice and their WT littermates fed 1% EPA-FFA, 1% DHA-FFA, 2.7% Lovaza or control diet were injected i.p. with AOM (7.5 mg/Kg) once weekly for 6 consecutive weeks to induce CRC. Mice were euthanized 20 weeks after the last AOM injection (n = 9-10 mice per group). (A, B) The gross colon images (A) and total colonic tumor number per mouse (B) for the indicated mouse groups are shown. (C-F) The levels of colonic 18-HEPE (C), colonic RvE1 (D), colonic 17-HDHA (E), and colonic RvD2 (F) per mouse for the indicated mouse groups, measured by LC/MS/MS, are shown. Lines for panels B-F represent means. Poisson regression for panel B, and two-way ANOVA with Bonferroni adjustments for all multiple comparisons for panels C-F. *\* P < 0.05;* and *\*\*\* P <0.001*.

### ALOX15 modulates the effects of EPA and DHA in EE and TG formulations on mutant *Apc*–induced CRC

To rigorously evaluate whether our findings were dependent on CRC model or the specific DHA and EPA formulations, we conducted a direct comparison of the effects of EPA-EE, EPA-TG, DHA-EE, and DHA-TG formulations, each as a 1% dietary supplement (Supplementary Table 5), on CRC in Apc^580mu^ mice and Apc^580mu^-ALOX15 mice. EPA and DHA in TG formulations were selected in this comparative testing because they have higher bioavailability than EE formulation (35). EPA-EE diet, but not EPA-TG, DHA-EE, or DHA-TG diet, significantly decreased colonic tumor multiplicity in Apc^580mu^ mice; Apc^580mu^-ALOX15-Gut mice had significantly lower colonic tumor multiplicity than Apc^580mu^ mice when fed DHA-EE or DHA-TG diet and trended to have lower colonic tumor multiplicity than Apc^580mu^ mice when fed control or EPA-TG diet (Fig. 3A and B). More importantly, ALOX15 expression reduced the multiplicity of large tumors (diameter > 2.5mm), which are associated with an increased invasiveness risk (36) (Fig. 3C). This trend was statistically significant for both DHA-EE and DHA-TG fed mice and was associated with fewer dysplastic lesions (Fig. 3C and D).

**Figure 3.**
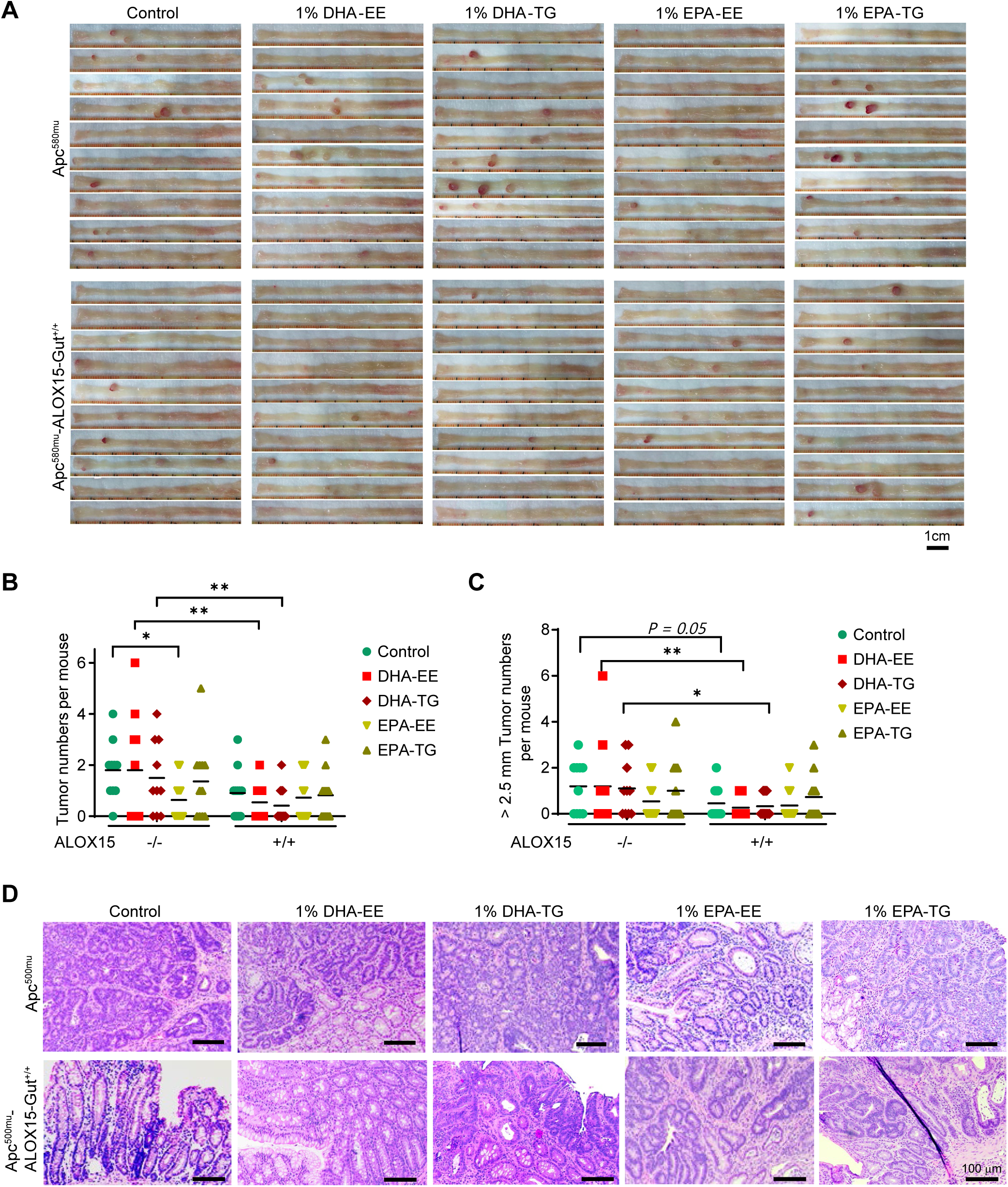
ALOX15 modulates the effects of DHA and EPA in EE and TG formulations on mutant Apc^580mu^–induced CRC. Apc^580mu^-ALOX15-Gut and Apc^580mu^ mice at 4-6 weeks were fed 1% DHA-EE diet, 1% DHA-TG diet, 1% EPA-EE diet, 1% EPA-TG diet or control diet for 10 weeks and then euthanized (n = 10-12 mice per group). (A-C) The gross colon images (A), total colonic tumor number per mouse (B), the multiplicity of the large tumors (diameter > 2.5mm) (C), and the representative H&E staining images of colon tumors (D) for the indicated mouse groups are shown. Lines represent means; Poisson regression for panels B and C.*\* P < 0.05;* and *\*\* P <0.01*.

Control diet–fed Apc^580mu^ mice had very low levels (0-1.33 ng/mg protein) of colonic 18-HEPE, RvE1, 17-HDHA, and RvD1-5. EPA-EE, EPA-TG, DHA-EE, or DHA-TG diet had no or minimal effects on these resolvins and their precursors in Apc^580mu^ mice (Fig. 4A-H, Supplementary Table 6). In contrast, EPA-EE, EPA-TG, DHA-EE, and DHA-TG diets in the presence of intestinal ALOX15 expression markedly increased resolvins E1, D2-5 and their precursors in Apc^580mu^-ALOX15-Gut mice (Fig. 4A-H, Supplementary Table 6). Intestinal ALOX15 expression increased 18-HEPE and RvE1 to higher levels in mice fed EPA-EE and EPA-TG than in the same mice fed DHA-EE and DHA-TG (Fig 4A-B, Supplementary Table 6). Similarly, intestinal ALOX15 expression increased 17-HDHA and RvD2-5 to higher levels in mice fed DHA-EE and DHA-TG than in the same mice fed EPA-EE or EPA-TG (Fig. 4C-H, Supplementary Table 6). Among the examined resolvins D1-5, colonic RvD1 level was lowest in Apc^580mu^ mice (0-0.56 ng/mg protein); ALOX15 significantly increased RvD1 levels in Apc^580mu^-ALOX15-Gut mice fed control diet, but not in the same mice fed EPA-EE, EPA-TG, DHA-EE or DHA-TG (Fig. 4D, Supplementary Table 6). RvD5 was the major product (mean concentration: 10.05-31.99 ng/mg protein), with mean concentration that was 10-folds higher than the other four RvDs (all mean concentrations: ≤ 3.00 ng/mg protein) in Apc^580mu^-ALOX15-Gut mice (Fig. 4H, Supplementary Table 6). Colonic levels of 18-HEPE, RvE1, 17-HDHA, RvD2-RvD5, but not RvD1 were significantly negatively correlated with colonic tumor multiplicity in Apc^580mu^ and Apc^580mu^ -ALOX15-Gut mice (Supplementary Fig. 3A-H). 17-HDHA and RvD5 were identified to be the main products differentially altered by intestinal ALOX15 transgenic expression with dietary DHA-EE or DHA-TG via volcano blot analyses (Fig. 4I, J).

**Figure 4.**
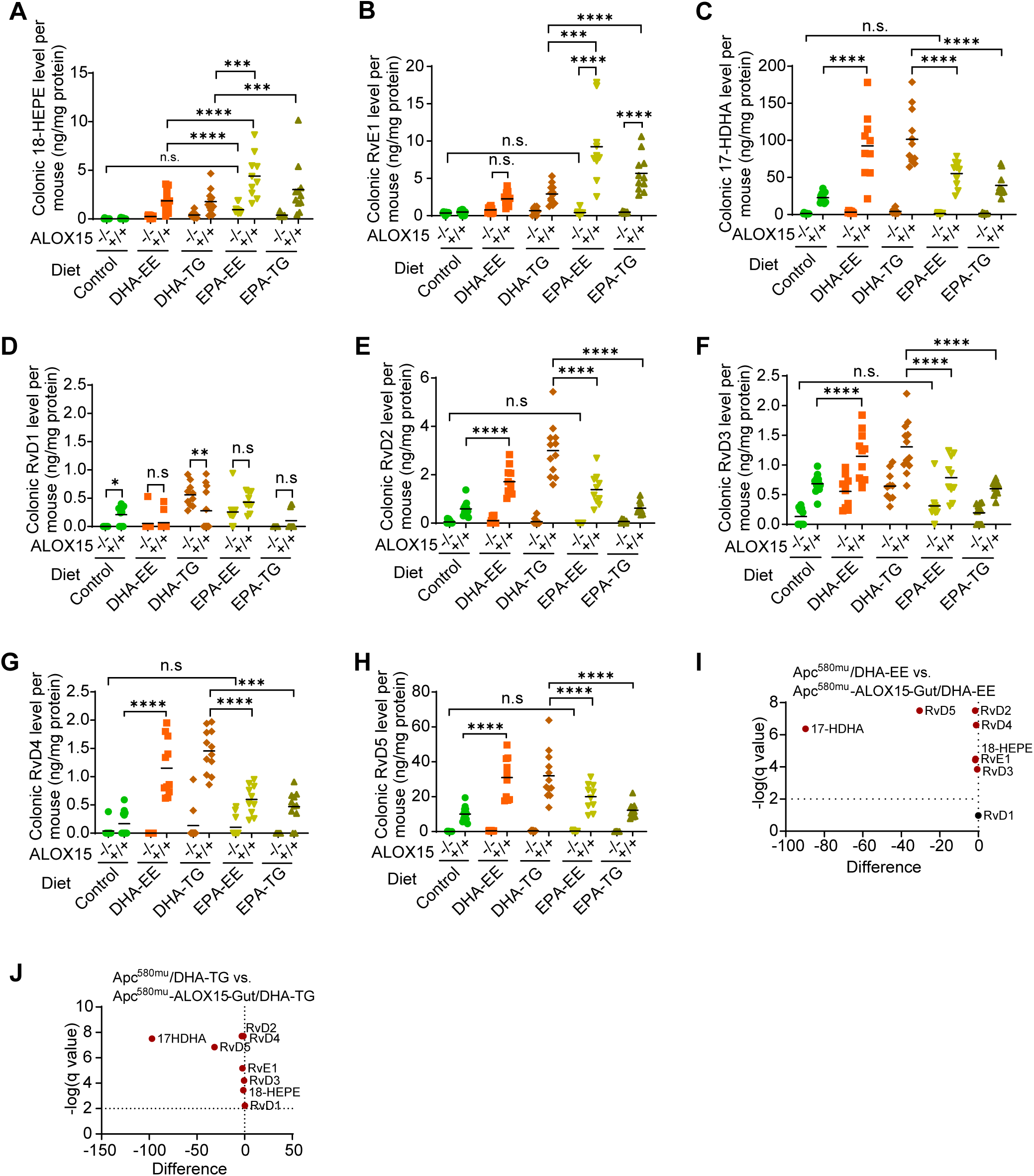
ALOX15 modulates the effects of DHA and EPA in EE and TG formulations on the generation of resolvins in Apc^580mu^ and Apc^580mu^-ALOX15 mice. Apc^580mu^-ALOX15-Gut and Apc^580mu^ mice were fed the diets described in Fig. 3 (n = 10-12 mice per group). (A-H) The levels of colonic18-HEPE (A), RvE1 (B), 17-HDHE (C) and RvD1-5 (D-H) in the scraped colonic mucosa cells per mouse from the indicated mouse groups, measured by LC/MS/MS, are shown. (I, J) Volcano plots of the differential alterations in the indicated colonic resolvin metabolites of DHA in the indicated mice. Comparisons of DHA-EE diet–fed Apc^580mu^ mice vs. DHA-EE diet–fed Apc^580mu^-ALOX15-Gut mice (I) and DHA-TG diet–fed Apc^580mu^ mice vs. DHA-TG–diet fed Apc^580mu^-ALOX15-Gut mice (J). Lines in panels represent means; two-way ANOVA with Bonferroni adjustments for all multiple comparisons for panels A-H. *\*\* P < 0.01; *** P < 0.001; **** P <0.0001*; and n.s.: non-significant.

### A clinically relevant dose of Lovaza and intestinal ALOX15 transgenic expression cooperate to inhibit AOM–induced CRC

Our initial experiments evaluated a commonly used dietary Lovaza diet (2.7%) tested in AOM (7.5mg/kg)–induced CRC in WT, ALOX15-Gut^+/-^, and ALOX15-Gut^+/+^ mice and showed dietary Lovaza was sufficient to inhibit CRC in WT mice; these effects were enhanced by ALOX15 in ALOX15-Gut^+/+^ mice (Fig. 2A, B). However, the highest FDA approved human Lovaza dose is equal to a 0.25% dietary Lovaza murine dose, which is much lower than the Lovaza doses commonly used in the published mouse studies. To simulate the human Lovaza usage dose, we therefore examined the effects of 0.25% dietary Lovaza on AOM–induced CRC in mice with or without intestinal ALOX15 transgenic expression. AOM induces CRC with AOM doses ranging from 7.5 mg/Kg to 10 mg/Kg, and the tumor burden is AOM dose dependent (37). In this study, we used high dose AOM (10mg/Kg) to increase the rigor of CRC testing given that the mice treated with 7.5mg/Kg AOM developed relatively low numbers of colorectal tumors (Fig. 2A, B; Supplementary Fig. 1A, B) (37). Moreover, in this study, we also investigated whether the effects of intestinal ALOX15 transgenic expression on CRC are dependent on the promoters that drive ALOX15 expression in mice. We therefore generated a new inducible ALOX15 mouse model (iALOX15) driven by Rosa26 constitutive promoter, in which ALOX15 protein was expressed and enzymatically active in colonic epithelial cells when breeding them with the mice that have intestinally directed Cre recombinase expression (e.g., Villin-Cre mice or CDX2-Cre mice) (Supplementary Fig. 4A-G). iALOX15; Villin-Cre mice generated by breeding iALOX15 mice with Villin-Cre mice were named iALOX15-Gut.

iALOX15-Gut mice had significantly lower colonic tumor volumes than Villin-Cre control mice when fed the control diet (Fig. 5A, B). Colonic tumor volumes trended to be smaller in WT mice fed 0.25% Lovaza diet than in the same mice fed control diet (P = 0.099); the Lovaza effects became stronger and statistically significant in iALOX15-Gut mice (Fig. 5A, B). More importantly, large colonic tumor (diameter > 3mm) multiplicity was only significantly reduced in iALOX15-Gut mice fed 0.25% Lovaza diet (Fig. 5A, C).

**Figure 5.**
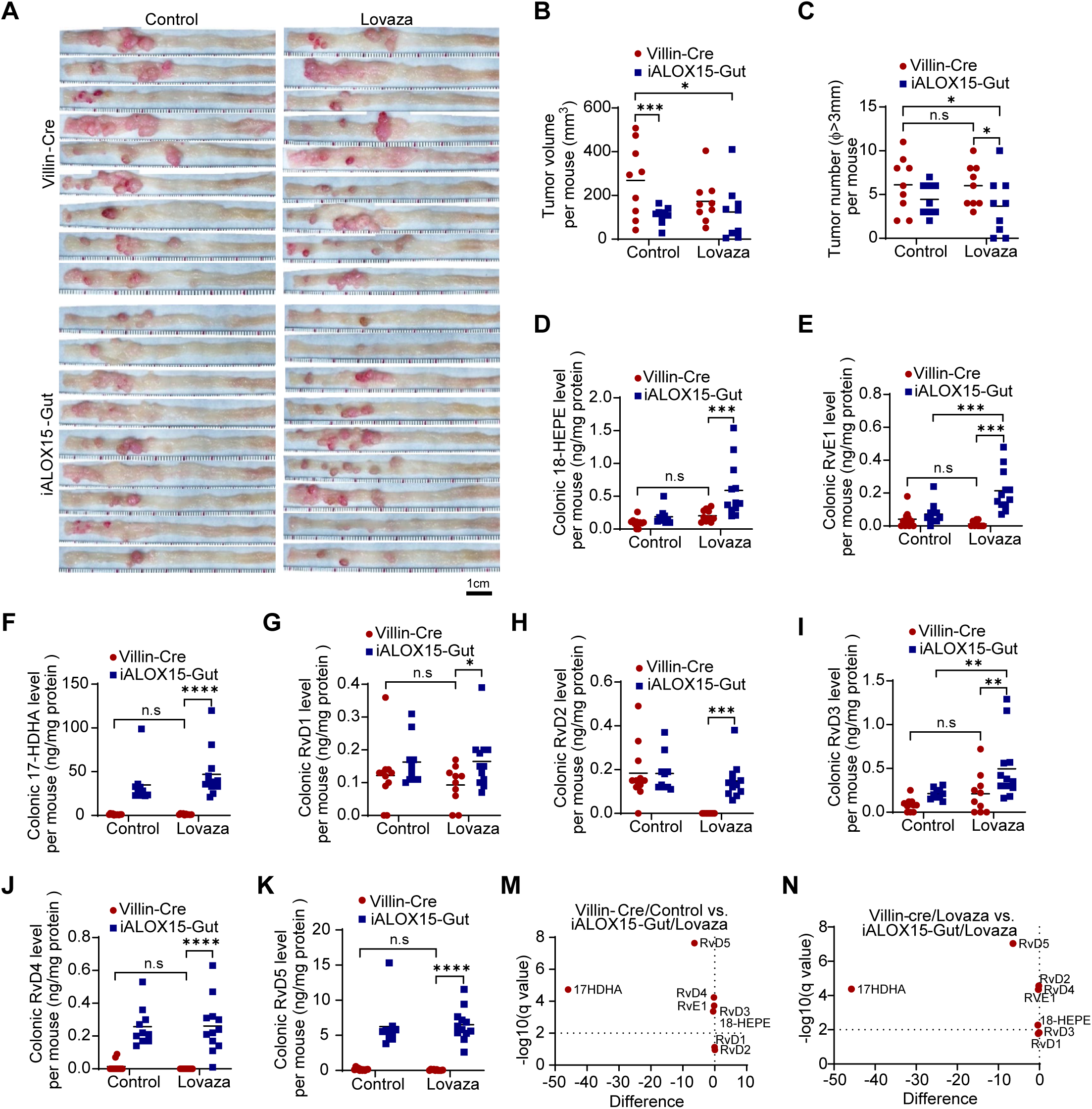
ALOX15 modulates the effects of clinically relevant Lovaza dose on AOM–induced CRC and the generation of resolvins using a novel inducible ALOX15 mouse model. 6-8 weeks iALOX15-Gut mice and their Villin-Cre control littermates fed 0.25% Lovaza diet or control diet were injected via i.p. with AOM (10mg/Kg) once weekly for 6 consecutive weeks to induce CRC. The mice were euthanized at 20 weeks after last AOM injection (n = 9 per group). (A-C) Gross colon images (A) and the colonic tumor volume (B) and large colonic tumor number (diameter > 3mm, C) per mouse for the indicated mouse groups are shown. (D-K) The levels of colonic18-HEPE (D), RvE1 (E), 17-HDHE (F) and RvD1-5 (G-K) in colonic crypts per mouse from the indicated mouse group, measured by LC/MS/MS, are shown. (M, N) Volcano plots of the differential alterations in the indicated resolvin metabolites of Lovaza in the indicated mice. Comparisons of control diet–fed Villin-Cre mice vs. Lovaza diet–fed iALOX15-Gut mice (M) and Lovaza diet–fed Villin-Cre mice vs. Lovaza diet–fed iALOX15-Gut mice (N). Lines in panels B-K represent means. Poisson regression for panel C, and two-way ANOVA with Bonferroni adjustments for all multiple comparisons for panels B, D-K. *\*P < 0.05; ** P <0.01; *** P < 0.001; **** P <0.0001*; and n.s.: non-significant.

The levels of colonic 18-HEPE, RvE1, 17-HDHA, and RvD1-5 were similar between control diet– and 0.25% Lovaza diet–fed Villin-Cre mice (Fig. 5D-K). In contrast, iALOX15-Gut mice fed 0.25% Lovaza diet had significantly higher levels of colonic 18-HEPE, RvE1, 17-HDHA, RvD1-5 than Villin-Cre mice fed the same diet (Fig. 5D-K). 17-HDHA and RvD5 were also the most differentially increased products in iALOX15-Gut mice fed 0.25% Lovaza diet (Fig. 5M, N). These data demonstrate that ALOX15 was required for this clinically relevant dose of Lovaza to generate resolvins and inhibit AOM–induced CRC; the effects of ALOX15 were independent of the promoters that drive ALOX15 expression.

### ALOX15 modulates the effects of EPA and DHA on the production of the chemokines and cytokines

During malignant transformation, colonic epithelial cells secret chemokines and cytokines to induce the formation of immune suppressive cells in the tumor microenvironment (TME) (e.g., tumor associated macrophages [TAMs]), thereby promoting CRC by suppressing anti-tumor immune response mechanisms such as recruitment of cytotoxic CD8 T cells (38). Thus, we investigated the effects of ALOX15 intestinal transgenic expression and subsequent resolvin formation on TAMs and effectors CD8 T cells abundance in TME in mice with AOM-induced CRC fed Lovaza or control diet. ALOX15 intestinal expression and formation of resolvins (Fig. 5) led to a reduction in M2-like TAMs (F4/80^+^/CD206^+^ cells) but an increase in CD8a^+^ T cells especially in TME of the mice fed Lovaza diet (Fig. 6A-C).

**Figure 6.**
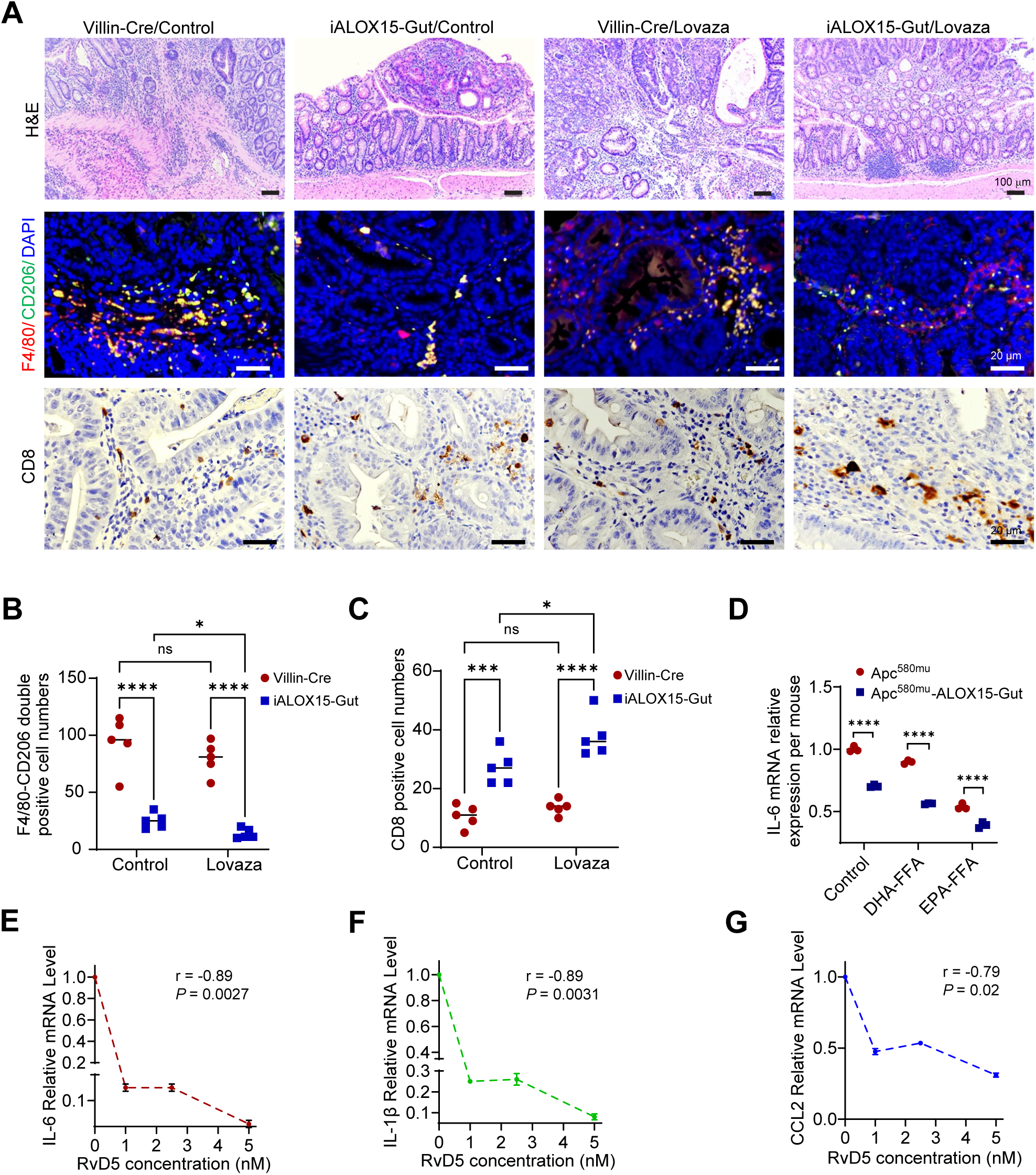
ALOX15 modulates the impact of Lovaza on CRC tumorigenesis by regulating the infiltration of tumor immune cells and their subsequent generation of pro-tumorigenic chemokines and cytokines. (A-C) The mice in the indicated groups as described in Fig. 5 were euthanized, and colonic tissues were harvested for further analyses (n=5 per group). (A) Representative images of H&E staining of colons (Top) and co-IF staining of CD206 with F4/80 (M2-TAMs) (Middle) and IHC staining of CD8a (Bottom). (B, C) The quantitative results of M2-TAMs (B) and CD8 T cells (C) for the indicated mice. (D) The levels of IL-6 mRNA expression in Raw 264.7 cells treated with organoid conditioned media from the indicated mice with the addition of 50ng/ml LPS. Values are means ± SEM for panels B-G. Two-way ANOVA with Bonferroni adjustments for all multiple comparisons for panels B-D. *\*P < 0.05; *** P < 0.001; **** P <0.0001*; and n.s.: non-significant. (E-G) Raw 264.7 cells treated with 0, 1, 2.5 and 5.0 nM RvD5 with 1ng/ml LPS were harvested and processed for total RNA extraction. The mRNA expression levels of IL-6 (E), IL-1β (F) and CCL2 (G) in Raw 264.7 cells measured by qRT-PCR are shown. Spearman correlation analyses of RvD5 concentrations with mRNA expression levels of IL-6 (E), IL-1β (F) and CCL2 (G) in Raw 264.7 cells.

To determine whether ALOX15 enzymatic metabolism of EPA-FFA and DHA-FFA in mice affected colonic production of chemokines and cytokines crucial for recruiting monocytes and polarizing them to TAMs, thus suppressing immune response to CRC (39–42), we screened 13-inflammatory chemokines in the scraped colonic mucosa cells of Apc^580mu^ mice and Apc^580mu^-ALOX15-Gut mice fed 1% EPA-FFA or 1% DHA-FFA or control diet for 6 weeks by using the LEGENDplex Mouse Proinflammatory Chemokine Panel kit (43). Intestinal ALOX15 expression decreased colonic CCL3-5 and CXCL5 levels in the mice fed control or DHA-FFA diet, but only CCL5 and CXCL5 were reduced when fed EPA-FFA diet (Supplementary Fig. 5A-D). To explore the potentially translational significance of these chemokines, we interrogated the publicly TCGA data available via UCSC-Xena portal and found that CCL3-5 and CXCL5 expression levels were significantly higher in human colon cancer tissues than their adjacent normal colon tissues (Supplementary Fig. 5E-H).

Cell–secreted resolvins can modulate cytokine production by neighboring cells such as macrophages including TAMs (44, 45). We therefore evaluated the effects of ALOX15 modulation of EPA-FFA and DHA-FFA metabolism in colonic epithelial cells on macrophage–secreted pro-tumorigenic IL-6 production using conditioned media from colonic organoids derived from Apc^580mu^ and Apc^580mu^-ALOX15-Gut mice fed 1% DHA-FFA or 1% EPA-FFA or control diet. Mouse macrophages, Raw 264.7 cells, incubated with organoid-conditioned media from EPA-FFA diet–fed Apc^580mu^ mice had lower IL-6 mRNA expression than those incubated with organoid-conditioned media from control diet*–*fed Apc^580mu^ mice; a parallel reduction in IL-6 mRNA expression was moderate when comparing DHA-FFA diet to control diet. Organoid-conditioned media from Apc^580mu^-ALOX15-Gut mice fed control, or EPA-FFA, or DHA-FFA diet significantly decreased IL-6 mRNA expression than the conditioned media of Apc^580mu^ mice fed the same diet (Fig. 6D). These findings suggested that ALOX15 expression in intestinal epithelial cells via its oxidative metabolism of EPA and DHA generated resolvins that were secreted into organoid-conditioned media from Apc^580mu^-ALOX15-Gut mice, which subsequently suppressed IL-6 production in macrophages. We therefore examined the effects of RvD5, the most abundant colonic resolvin formed by intestinal ALOX15 transgenic expression in the mice (Fig. 4, 5), on lipopolysaccharides (LPS)-induced IL-6, IL-1β and CCL2 expression in Raw 264.7 (Supplementary Fig. 6A-C). RvD5 treatment inhibited LPS–induced IL-6, IL-1β and CCL2 expression in Raw 264.7 cells in a dose dependent manner (Fig. 6F-G). These findings demonstrated that RvD5, which was abundantly produced from DHA by ALOX15 expression in colonic epithelial cells (Fig. 4, 5), inhibited the macrophage production of important pro-tumorigenic cytokines (e.g., IL-6, IL-1β and CCL2) as a potential mechanism for CRC suppression by EPA and DHA in the presence of intestinal ALOX15 expression.

## DISCUSSION

Our findings demonstrate that the effects of EPA and DHA on CRC were divergent and occasionally altered by their different formulations, but more importantly, by intestinal expression of ALOX15, a host factor that regulates production of resolvins from EPA and DHA.

Fish oil and its derivatives have variable effects on CRC. Contrary to previous reports showing that dietary fish oil reproducibly suppresses CRC in preclinical rodent studies (6–8, 34), we found that fish oil promoted AOM/DSS-induced CAC in mice. These discrepant results are unlikely to be attributed to differences in fish oil preparations as our study employed fish oil dietary supplement similar to those used in prior contradictory studies (34). Other reports have also shown that fish oil, particularly when enriched with DHA, promotes CAC (9, 10, 46), which is consistent with our findings. In contrast, we found that EPA-FFA inhibited CAC in the same experimental model. These data, along with the studies from other investigators, indicate that fish oil and its derivatives have variable effects on CRC. Identification of factors that determine the effects of fish oil and its derivative on CRC is crucial to guide the appropriate use of the widely consumed dietary EPA and DHA supplements and FDA approved treatments.

EPA and DHA differentially modulate CRC. This notion has been previously postulated mainly based on indirect comparisons of various EPA and DHA formulations in preclinical studies using CRC animal models that have limited translational applications to human CRC. For examples, EPA-FFA inhibited AOM/DSS–induced CAC and mutant *Apc–*induced intestinal tumors in Apc^min^ mice (11, 12), while DHA– enriched fish oil promoted CAC in AOM/DSS–treated mice (9, 10, 46). In one published study that directly compared the effects of EPA-EE and DHA-EE on CRC, EPA-EE decreased while DHA-EE increased colonic tumor multiplicity in Apc^min^ mice (47). Apc^min^ mice however mainly form tumors in the small intestine whereas most human intestinal tumors occur in the large intestine. CAC mouse models are limited in representing human CRC as the vast majority of human CRC is unrelated to CAC. In our studies, we used the two most currently representative mouse models of human CRC in which CRC was induced either by the carcinogen AOM or intestinally-targeted *Apc* mutation via CDX-2 promoter–driven Cre recombinase expression (48, 49). Our direct comparisons evaluating the effects of different EPA and DHA formulations on CRC showed that 1) dietary EPA-FFA and EPA-EE inhibited CRC while DHA-FFA and DHA-EE failed to inhibit CRC, demonstrating that EPA and DHA in FFA and EE formulations had different effects on CRC; 2) the combination of EPA-EE and DHA-EE in Lovaza suppressed CRC, suggesting that DHA-EE effects on CRC were modified when combined with EPA-EE; 3) dietary EPA and DHA in the TG formulation that reportedly increased their bioavailability (50) failed to suppress CRC; and 4) intestinal ALOX15 transgenic expression significantly inhibited CRC in the mice fed DHA-FFA, DHA-EE, DHA-TG, and EPA-TG, and enhanced EPA-FFA suppression of CRC. Thus, our results collectively demonstrate that EPA and DHA, in three distinct formulations (FFA, EE, TG) administered individually or in combination, exhibited varied effects on CRC, and CRC suppressive effects were not enhanced with TG formulation with higher bioavailability. In contrast, intestinally expressed ALOX15 modulated the variable effects of EPA and DHA on CRC to consistently suppress CRC.

Intestinal ALOX15 transgenic expression is essential to colonic resolvin production from EPA and DHA. Our data show that intestinal ALOX15 expression in mice was critical for generation of both RvE and RvD resolvin series and their precursors from EPA and DHA. EPA and DHA supplementation without intestinal ALOX15 expression had minimal to no capacity in generating these metabolites. The role of ALOX15 in generating RvDs has been previously reported (51). While an ALOX15-like function of aspirin– acetylated COX-2 has been reported to mediate the generation of the RvEs’ precursor 18-HEPE from EPA (24), the direct role of ALOX15 in 18-HEPE formation is understudied. Our results showed that intestinal ALOX15 expression induced production of 18-HEPE from EPA in several in-vivo mouse models in the absence of aspirin-acetylated COX2. These findings demonstrate the direct role of ALOX15 in oxidative metabolism of EPA into18-HEPE. Moreover, our findings revealed, in multiple CRC mouse models, the critical regulatory role of ALOX15 in the generation of RvE1, 17-HDHA, and RvD2-5 from diverse formulations of EPA and DHA administered individually or in combination (e.g., Lovaza), including a Lovaza dose that simulates the FDA approved human Lovaza dose.

Colonic resolvins, induced by intestinal ALOX15 expression, play a critical role in suppressing CRC. The levels of colonic RvE1, RvD 2-5 and their precursors (18-HEPE and17-HDHA) were negatively correlated with colonic tumor multiplicity in the mice, suggesting their contributions to CRC suppression. Dietary supplementation with DHA-FFA, DHA-EE, DHA-TG, EPA-TG, and a clinically relevant dose of Lovaza (0.25%) resulted in consistent CRC suppression only when combined with intestinal ALOX15 transgenic expression that induced the generation of colonic resolvins. Furthermore, a clinically relevant dose of Lovaza also required intestinal ALOX15 expression to effectively suppress large colorectal tumor formation. These findings underscored the pivotal role of ALOX15 as a host factor that drives the generation of resolvins from EPA and DHA to subsequently suppress CRC.

Resolvins suppress important pro-tumorigenic chemokines and cytokines as a mechanism to inhibit CRC. Intestinal barrier integrity is compromised during the early phases of CRC, which allows bacterial products—especially LPS, the most abundant endotoxin generated by intestinal dysbiosis—to enter the colonic TME and trigger the production of chemokines and cytokines, and subsequently, recruitment of TAMs (39–42). TAMs, which compose up to 50% of the TME, generate additional pro-tumorigenic chemokines and cytokines (e.g. IL-6, CCL2) to expand the TAM population and recruit other immunosuppressive myeloid cell populations into the TME, leading to CRC progression (19, 20, 52–56). We found that intestinal ALOX15 expression and subsequent production of colonic resolvins in mice repressed TAM recruitment while increasing the abundance of effector CD8 cells. Previous studies showed that supplementation of exogenous resolvins suppressed the growth of grafted murine isogenic cancer cells in mice via modulating immune cells within TME (21, 57, 58). Importantly, our findings provide first evidence that endogenously generated resolvins, via specific ALOX15 expression in colonic epithelial cells, alleviated TME immune suppression in an autochthonous CRC mouse model.

Colonic epithelial cells with ALOX15 expression that generated resolvins repressed the production of CCL3-5 and CXCL5 by colonic epithelial cells to recruit TAMs. These findings are pertinent for understanding the mechanisms by which resolvins impact human CRC. CCL3-5 and CXCL5 are upregulated in human colorectal cancers as we have found through analysis of publicly available databases. These data are in agreement with prior reports showing CCL3 and CCL4 are upregulated in colorectal cancers; CCL4 increases the recruitment of TAMs into TME (59); CCL5 contributes to CRC progression (60, 61); and CXCL-5 recruits myeloid suppressive cells into TME (62).

ALOX15 expression in colonic epithelial cells not only affects chemokines and cytokines production in colonic epithelial cells but also in macrophages. This concept is supported by our findings showing conditioned media of organoids derived from colonic epithelial cells of mice expressing ALOX15 expression fed EPA-FFA or DHA-FFA diet significantly repressed IL-6 expression in macrophages compared to those from WT mice fed the same diet. Furthermore, RvD5, the predominant resolvin induced by intestinal ALOX15 expression, directly inhibited the expression of LPS-induced IL-6, IL-1β, and CCL-2 in mouse macrophages. IL-6, IL-1β, CCL2 are known to promote CRC progression (63–65). Therefore, intestinal ALOX15 expression by increasing the production of RvD5 and other resolvins from EPA and DHA inhibited macrophage production of pro-tumorigenic cytokines and subsequently suppressed CRC.

In summary, our collective findings demonstrate that intestinal ALOX15 expression is required for generating resolvins from EPA and DHA, leading to reduced expression of pro-tumorigenic cytokines and chemokines by colonic tumors cells and M2-like TAMs, subsequently suppressing CRC (Fig. 7). Consequently, ALOX15 emerges as a crucial host factor that should be considered in the development of strategies utilizing EPA and DHA for CRC suppression.

**Figure 7.**
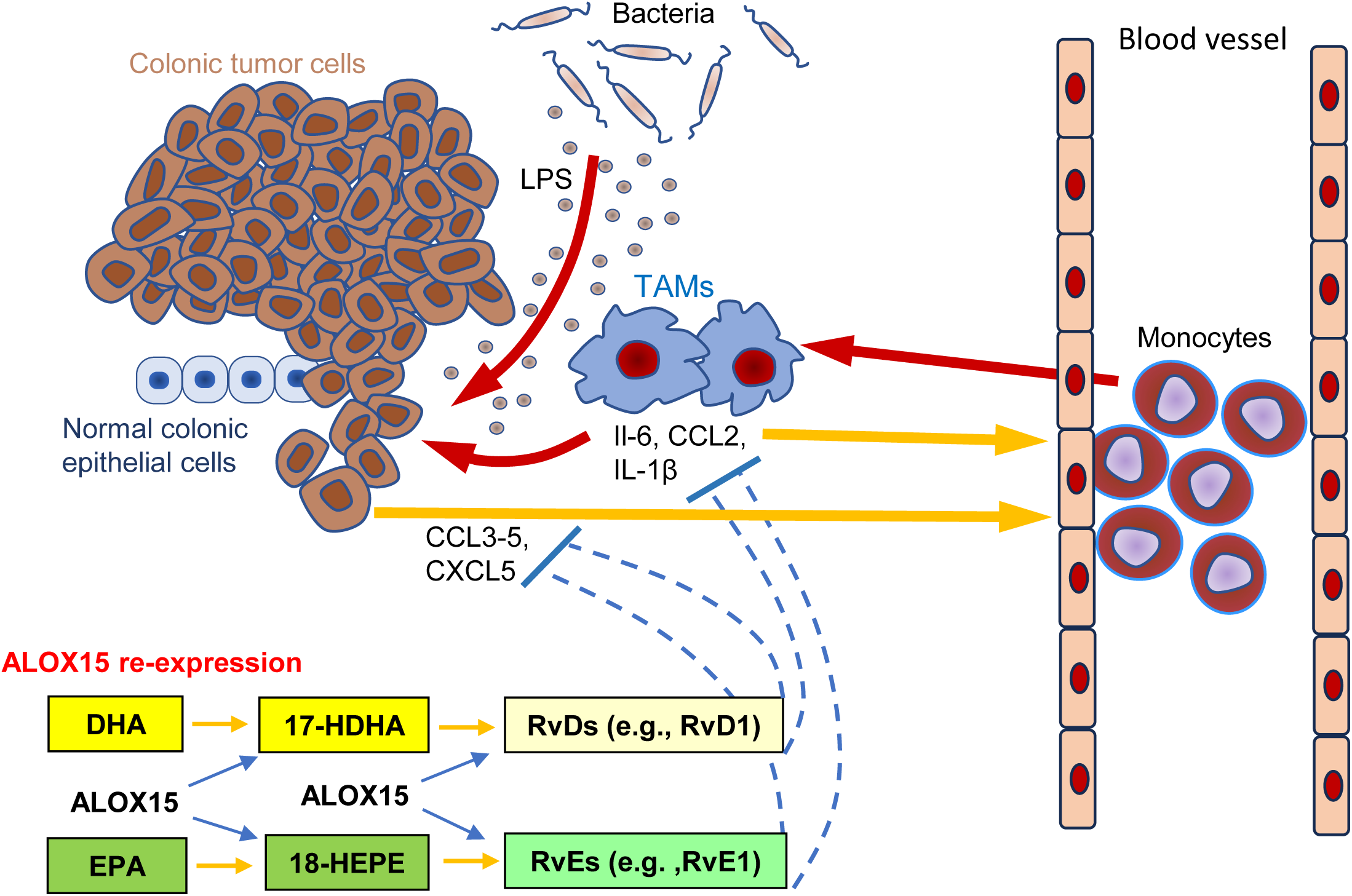
Conceptual model of this study: ALOX15-generated resolvins inhibit the productions of chemokines and cytokines by colonic tumor cells and M2-like TAMs, consequently suppressing CRC.

## METHODS

### Cell culture

RAW 264.7 mouse macrophages were purchased from ATCC (TIB-71^™^), and the RAW 264.7 cells were cultured and expanded in Dulbecco modified Eagle medium (high glucose) supplemented with 10% fetal bovine serum (FBS) and 1% penicillin/streptomycin, and cultured in humidified atmosphere with 5% CO_2_ at 37°C.

### Animals

Mouse care and experimental protocols were approved by and conducted in accordance with the guidelines of the Animal Care and Use Committee of The University of Texas MD Anderson Cancer Center. We generated the mice with targeted ALOX15 overexpression in intestinal epithelial cells driven by a Villin promoter (designated as ALOX15-Gut^+/-^ mice) as described previously (66). Moreover, we also worked with PolyGene Transgenetics, Switzerland, to generate a novel ALOX15 chimera mouse model with an inducible/conditional ALOX15 expression at the Rosa26 locus through a 4-step procedure: target vector construction, embryonic stem cell culture, blastocyst injection, and Flp-/Germ line breeding (designated as iALOX15^+/-^ mice). Refer to Supplementary Fig. 4 for the details of this mouse generation and characterization. To further generate the iALOX15 chimera mice with the genetic FVB/N background that is suitable to the AOM-induced CRC model, we backcrossed the iALOX15^+/-^ chimera mice driven by the Rosa promoter with FVB/N WT mice for at least 10 generations. FVB/N genetic background (>95%) was confirmed by genetic background characterization with a set of 80-100 informative markers (SNPs and/or SSLPs) by the MD Anderson Laboratory Animal Genetic Services Core Facility. *B6.Cg-Tg (*CDX2*-Cre) 101Erf/J* (CDX2-Cre, #009350) mice were purchased from Jackson Laboratory. Villin-Cre mice with an FVB/N genetic background were a gift from Dr. Sue Fischer as described previously (67). Apc^Δ580^-flox mice, in which the *Apc* exon 14 is flanked with loxP sites, were a gift from Dr. Kenneth E. Hung at Tufts Medical Center (68). Apc^Δ580^-flox mice were bred with CDX2-Cre mice to generate Apc^Δ580^-flox^+/-^; CDX2-Cre^+/-^ mice (designated as Apc^580mu^ mice), in which exon 14 of the *Apc* gene was deleted to consequently generate a codon 580 frame-shift mutation in colonic epithelial cells (69).

### Generation of experimental mice

1. ALOX15-Gut^+/-^ mice were bred with ALOX15-Gut^+/-^ mice to generate ALOX15-Gut^+/+^ mice (designated as ALOX15-Gut in this manuscript, unless specified), in which homozygous ALOX15 was expressed specifically in Villin-expressing intestinal epithelial cells, and control wild type (WT) littermates.
2. Apc^580mu^; ALOX15-Gut^+/-^ mice were bred with ALOX15-Gut^+/-^ mice to generate Apc^580mu^; ALOX15-Gut^+/+^ mice (designated as Apc^580mu^-ALOX15-Gut) and Apc^580mu^ littermates, in which CDX2-Cre–driven colonic Apc genetic mutation and a Villin-promoter–driven intestinal homozygous ALOX15 overexpression or CDX2-Cre–driven colonic Apc genetic mutation only, were induced, respectively.
3. iALOX15^+/-^ mice; Villin-Cre^+/-^ were bred with iALOX15^+/-^ mice to generate iALOX15^+/+^; Villin-Cre^+/-^ (designated as iALOX15-Gut), in which homozygous ALOX15 was expressed specifically in Villin-expressing intestinal epithelial cells, and Villin-Cre^+/-^ (designated as Villin-Cre) littermates.

### CRC gross evaluation in multiple mouse models fed diets supplemented with EPA and DHA in the three different formulations (FFA, EE and TG)

1. The effects of fish oil or EPA-FFA on AOM/DSS–induced CAC in ALOX15-Gut and WT mice. 6-8 weeks-old ALOX15-Gut mice and their WT littermates were fed with control diet containing 15% corn oil (D12070201, Research Diets), and fish oil diet containing 11.5% menhaden oil and 3.5% corn oil (D12070202, Research Diets) (the details for these customized diets are described in Supplementary Table 1); or fed control diet containing 7% corn oil (D03090904P, Research Diets) and EPA-FFA diet containing 1% EPA-FFA, provided as an unrestricted gift from SLA Pharma AG (Switzerland) (https://www.slapharma.com/) and 6% corn oil (D12090801, Research Diets) (the details for these customized diets were described in Supplementary Table 3) for 4 weeks before the mice were administrated with 10 mg/kg AOM via intraperitoneal (i.p) injection. One week after the AOM injection, the mice were treated with 1.5% DSS water for 7 days and then regular water for 14 days as a cycle, and this cycle was repeated for 3 consecutive cycles. The mice continued to feed the assigned diets when treated with AOM and DSS.
2. The effects of EPA-FFA, DHA-FFA, and Lovaza on 7.5mg/Kg AOM–induced CRC in ALOX15-Gut and WT mice. 6-8 weeks-old ALOX15-Gut mice and their WT littermates were fed with customized diets (diet details described in Supplementary Table 3): either 7% corn oil control diet (D03090904P, Research Diets), 1% EPA-FFA diet (D12090801, Research Diets), DHA-FFA diet containing 1% DHA-FFA (purity: ≥98.0%, catalog# 90310, Cayman Chemical) and 6% corn oil (D12090806, Research Diets), or Lovaza diet containing 2.7% Lovaza (Woodward pharma) and 4.3% corn oil (D12090803, Research Diets) for 4 weeks before the mice were administrated with 7.5mg/kg AOM via i.p. injection once a week for 6 consecutive cycles while the mice continued to feed the assigned diets. The mice were euthanized 20 weeks after the last AOM injection.
3. The effects of EPA-EE, EPA-TG, DHA-EE and DHA-TG on Apc^580mu^-induced CRC in ALOX15-Gut and WT mice. 4-6 weeks-old Apc^58mu^ and Apc^580mu^-ALOX15-Gut mice were fed 7% corn oil control diet (D03090904P, Research Diets), 1% DHA-FFA (D12090806, Research Diets) (diet details described in Supplementary Table 3); or fed 7% corn oil control diet (TD.120422, Envigo), 1% EPA-EE diet (EPA-EE: catalog# U-99-E, NU-CHEK PREP, INC; the diet: TD.160449, Envigo), 1% EPA-TG diet (EPA-TG: catalog# T-325, NU-CHEK PREP, INC; the diet: TD.160151, Envigo), 1% DHA-EE diet (DHA-EE: catalog# U-84-E, NU-CHEK PREP, INC; the diet: TD.160450, Envigo), or 1% DHA-TG diet (DHA-TG: catalog# T-310, NU-CHEK PREP, INC; the diet: TD.160452, Envigo) (diet details described in Supplementary Table 6) for 10 weeks.
4. The effects of 0.25% Lovaza on 10mg/Kg AOM-induced CRC in iALOX15-Gut and WT mice. 6-8 weeks-old iALOX15-Gut mice and their WT littermates were fed either 7% corn oil control diet (D03090904P, Research Diets) or 0.25% Lovaza diet (D20042001, Research Diets) for 4 weeks before the mice were administrated 10mg/kg AOM via i.p. injection once a week for 6 consecutive cycles while the mice continued the assigned diets. The mice were euthanized 20 weeks after the last AOM injection.

The mice from all four experiments described above were then euthanized. The colons from the rectum to the cecum were removed and washed with phosphate-buffered saline (PBS) and photographed, tumors were counted, and tumor sizes were measured. The tumor volumes were calculated using the formula V (volume) = (length ×width^2^)/2 (70). The distal one-third of colons were collected in 10% neutral formalin for hematoxylin and eosin staining and the other two-thirds of colons were scraped to harvest colonic epithelial cells for further analyses (e.g., RNA, protein, and mass spectrometry), as described in the corresponding supplementary method sections.

### Mouse colonic tissue histology

Formalin-fixed samples were embedded in paraffin and sectioned onto slides at 5 μm thick and then stained with H&E. Digital HE staining slides for mouse colonic tissues were scanned with Aperio AT2 (Leica biosystems) and the scanned images were captured with Aperio ImageScope software [v12.3.3.5048].

### Measurement of EPA and DHA in mouse sera and mouse colonic mucosa cells and by Liquid chromatography and tandem mass spectrometry (LC-MS/MS)

Sample preparation: 1) 10 μl of serum sample was diluted with 20 μl of 5 N HCl containing 10 μl of internal standards (EPA-d_5_ and DHA-d_5_, 10 μg/ml for each). 400 μl of acetonitrile was then added to this solution and the mixture was hydrolyzed at 100°C for 1 hour. After hydrolysis, 1 ml of chloroform and 1 ml of water were added followed by vortex mixing for 1 min, and then centrifuged at 3000 rpm at 4°C for 10 min. The organic layer was collected, pooled and evaporated to dryness under a stream of nitrogen at room temperature. The sample was then reconstituted in 100 μl of methanol. 2) Mouse mucosa cell pellets were directly mixed with 20 μl of 5 N HCl containing 10 μl of internal standards (EPA-d_5_ and DHA-d_5_, each for 10 μg/ml) and 400 μl of acetonitrile and then sonicated for 3 min on ice. 300 μl of homogenate was transferred and subjected to hydrolysis step and the extraction procedure was followed as described for serum sample.

The LC-MS/MS analysis was performed using Agilent 6460 Triple Quad mass spectrometer equipped with an Agilent 1200 HPLC system (Agilent Technologies). EPA and DHA were separated using Luna C_5_ column (5 µm, 2.0 x 50 mm) (Phenomenex, Torrance, CA) by isocratic elution. The mobile phase consisted of 10 mM ammonium acetate in water (phase A, 15%) and methanol (phase B, 85%). Flow rate was 300 μl/min with column temperature maintained at 25 °C. Sample Injection volume was 2 µl. EPA and DHA were detected using electrospray negative ionization and multiple-reaction monitoring (MRM) monitoring the transitions at m/z 301.2→257 for EPA, m/z 327.2→283 for DHA, m/z 306.2→262 for EPA-d_5_, and m/z 332.2→288 for DHA-d5. The mass spectrometer was operated in the electrospray negative ion mode with a gas temperature of 300°C, a gas flow rate of 10 ml/min, and a nebulizer pressure of 20 psi. The temperature of the sheath gas was 300°C, and the sheath gas flow rate was 12 l/min. The capillary voltage was -2900 V. The results were expressed as nanograms of EPA and DHA per ml for serum or nanograms per mg protein for colonic mucosa cells.

### Measurement of resolvins in mouse colonic mucosa cells by LC-MS/MS

Sample preparation: 1) mouse colonic mucosa cell pellets were homogenized in 300 μl of PBS containing 0.1% EDTA. 10 μl of 10% BHT and 10 μl of internal standard (100 ng/ml, PGE_2_-D4, Cayman Chemical) were added to 200 μl of homogenate, followed by addition of 1 ml of methanol. The mixture was vortexed and centrifuged at 14,800 rpm for 10 min at 4°C. The supernatant was collected and evaporated under a stream of nitrogen and reconstituted in 100 μl of methanol/0.1% acetic acid (1:1) which was then subjected to LC-MS/MS analysis. For the mouse serum sample, a similar extraction procedure as described above was followed except 100 μl of plasma and 800 μl of methanol being used.

Resolvins were measured by LC-MS/MS method. Briefly, analyses were performed using an Agilent 6460 Triple Quad mass spectrometer (Santa Clara, CA) equipped with an Agilent 1200 HPLC system. Resolvins were separated using an Agilent XDB C18 1.8-µm, 2.1 × 50-mm column. The mobile phases consisted of 0.1% formic acid in 10 mM ammonium acetate in water (phase A) and 0.1% formic acid in methanol (phase B). For the analysis of 18-HEPE, RvE1, 17-HDHA, RvD1, RvD2, RvD3, RvD4, and RvD5, the separation was achieved using a linear gradient of 20–90% of phase B with a total time of 20 min. The flow rate was 300 μl/min with a column temperature of 30°C. The sample injection volume was 15 μl. Samples were kept at 4 °C during the analysis. The mass spectrometer was operated in the electrospray negative ion mode with a gas temperature of 350°C, a gas flow rate of 10 ml/min, and a nebulizer pressure of 20 psi. The temperature of the sheath gas was 350 °C, and the sheath gas flow rate was 12 l/min. The capillary voltage was -2900 V. Fragmentation was performed for all compounds, using nitrogen as the collision gas. All resolvins were detected using electrospray negative ionization and MRM mode of the transitions at mass-to-charge ratios of 375.1→215.1 for RvD1, 375.2→174.9 for RvD2, 375.0→147.0 for RvD3, 375.0→101.0 for RvD4, 359.0→199.0 for RvD5, 349.1→195.0 for RvE1, 343.1→201.1 for 17-HDHA, and 317.2→215.1 for 18-HEPE. The results were expressed as nanograms of resolvins per milligram of protein for colonic mucosa cells.

### Quantification of LEGENDplex mouse proinflammatory chemokine panel in mouse colonic epithelial cells

Six-week-old Apc^580mu^-ALOX15-Gut mice and Apc^580mu^ littermates (n = 2-6 mice per group) fed either control or 1% DHA-FFA or 1% EPA-FFA or control diet for 6 weeks were euthanized, and their colons were scraped to harvest the colonic mucosa cells. The whole protein lysates of the harvested colonic mucosa cells were mechanically homogenized in protein lysis buffer (0.5% Nonidet P-40, 20 mM 3-[N-morpholino] propanesulfonic acid [pH 7.0], 2 mM ethylene glycol tetraacetic acid, 5 mM ethylenediaminetetraacetic acid, 30 mM sodium fluoride, 40 mM β-glycerophosphate, 2 mM sodium orthovanadate, 1 mM phenylmethylsulfonyl fluoride, and 1× complete protease inhibitor cocktail [Roche Applied Science]), and the protein concentrations were measured. A bead-based assay (LEGENDplex Mouse Proinflammatory Chemokine Panel, #740007; BioLegend, San Diego, CA) that uses the principles of sandwich enzyme-linked immunosorbent assay to quantify soluble analytes using a flow cytometer was used to quantify the chemokine concentrations (CCL2, CCL5, CXCL10, CCL11, CCL17, CCL3, CCL4, CXCL9, CCL20, CXCL5, CXCL1, CXCL13, and CCL22) per the manufacturer’s instructions. The data were collected and analyzed on BD FACS Canto II analyzer with FACSDiva software version 8.0 (BD) and analyzed using the LEGENDplex version 8.0 Data Analysis Software (BioLegend). The results for protein lysates were normalized to their corresponding protein and presented as pg/mg of protein, as described previously (71).

### Primary mouse colonic organoid culture and conditioned medium collection

Six-week-old Apc^580mu^-ALOX15-Gut mice and Apc^580mu^ littermates (n = 3 mice per group) fed either control or 1% DHA-FFA or 1% EPA-FFA or control diet for 6 weeks were euthanized, and their colons were harvested. The harvested colon tissues were digested with 10 mM ethylenediaminetetraacetic acid (EDTA) in chelation buffer (5.6 mM Na_2_HPO_4_, 8.0 mM KH_2_PO_4_, 96.2 mM NaCl, 1.6 mM KCl, 43.4 mM sucrose, 54.9 mM D-sorbitol, 0.5 mM DL-dithiothreitol) at room temperature for 10 min. Crypts were isolated by vigorously shaking the digested tissues for 30s and then allowing sedimentation for 1 min prior to collecting the supernatant. The isolated crypts were counted using a hemocytometer and embedded in growth factor reduced Matrigel (Cat# 356231, Corning) at 20 crypts/µL Matrigel. Approximately 20-μL droplets of Matrigel with crypts were seeded onto a flat-bottom 24-well low attachment plate (Sigma). The Matrigel was solidified for 15 min in a 37°C incubator, and then 250 µL/well organoid culture medium (advanced Dulbecco’s modified Eagle medium/F12 supplemented with penicillin/streptomycin, 2 mM GlutaMAX, 10 mM HEPES [all from Invitrogen], 100 ng/mL mouse recombinant Wnt-3A [Millipore], 50 ng/mL mouse epidermal growth factor [Invitrogen], 100 ng/mL mouse recombinant noggin [Peprotech], 1 µg/mL human R-spondin-1 [Peprotech], 1 mM N-acetyl-L-cysteine [Sigma-Aldrich], 1× N-2 [Invitrogen], 1× B-27 [Invitrogen]), and 10 µM Y-27632 (Sigma) for 72 hours, and the conditioned media were harvested for further use.

### Immunohistochemistry (IHC) and immunofluorescence (IF) staining

Tissue sections 5 µm thick were deparaffinized and rehydrated. Antigen retrieval was then performed by immersion of slides into Antigen Unmasking Solution (#H3300, Vector Laboratories) and heating in a steam chamber for 35 min. For IHC staining, slides were treated with 3% H_2_O_2_ solution to reduce endogenous peroxidase, incubated with blocking buffer (5% goat serum in TBST) for 30 min, and then incubated with primary antibodies overnight. The following primary antibodies were used: CD8a (#98941, 1:300) from Cell Signaling Technology. On the second day, the tissue sections were incubated with biotinylated secondary antibodies (VECTASTAIN ABC kit; Vector Laboratories) for 1 h, followed by incubation with avidin-coupled peroxidase (Vector Laboratories) for 30 min. The slides were developed using 3,3ʹ-diaminobenzidine (DAB) (Agilent Dako) and then counterstained with Mayer’s hematoxylin (Agilent Dako). For IF staining, the slides were incubated with blocking buffer (1.5% goat serum, 0.3% Triton X-100 in phosphate-buffered saline [PBS]) for 1 h at room temperature and then underwent primary incubation at 4°C overnight. The primary antibodies were: F4/80 (#71299,1:100) and CD206 (#24595,1:800) from Cell Signaling Technology. On the second day, the slides were incubated with the Alexa Fluor 488 or Alexa Fluor 594 fluorescence-conjugated secondary antibody at room temperature for 1 h, and then washed and mounted with ProLong Gold Antifade Mountant with DAPI (Thermo Fisher Scientific). The positive staining cells were counted under microscope with 20X magnification for 5 randomly selected fields, then averaged and presented as quantitative data for each mouse.

### RAW 264.7 mouse macrophage treatment and mRNA expression measurement of the cytokines

264.7 mouse macrophages were split in 6 wells of plates and cultured with harvested conditioned medium (described in previous section) mixed with fresh DMEM medium (1:1) with added LPS (50 ng/ml) for 24 hours or treated with 0, 1.0, 2.5 and 5.0 nM RvD5 (Cayman Chemical, catalog # 10007280) with added LPS (1 ng/ml) in 10% FBS of DMEM medium for 4 hours. The cells were then harvested in TRIzol (Invitrogen) for total RNA extraction according to the manufacturer’s protocol. cDNA was synthesized from 1µg total RNA per sample using a Bio-Rad cDNA Synthesis Kit (Bio-Rad Laboratory). Quantitative real-time polymerase chain reaction (RT-qPCR) analyses were performed using a FastStart universal probe master (Roche) and StepOnePlus PCR system (Applied Biosystems). All the qRT-PCR probes (mouse IL-6, IL-1β and CCL2) were purchased from Applied Biosystems. The relative RNA expression levels were normalized to the expression of mouse HPRT (Applied Biosystems) and calculated using a comparative threshold cycle method (ddC_t_) (71).

### Statistical methods

Statistical significance was determined by the unpaired Student *t*-test, chi-square test, or analysis of variance (one-way or two-way) with Bonferroni adjustments for all multiple comparisons as indicated for the various experiments. Tumor count/multiplicity analyses were performed using Poisson regression (67). The data were log-transformed as needed to accommodate the assumptions of normality and homoscedasticity implicit in the statistical methodologies employed. The significance of correlations was determined by Spearman correlation coefficient test. All tests were two-sided, and significance was defined as *P* < 0.05. Data were analyzed using SAS software, version 9.4 (SAS Institute) or GraphPad Prism 7.01 (GraphPad Software). The data are presented as mean ± standard deviation or mean ± standard error as indicated in the corresponding legends (\**P* < 0.05, \*\**P*<0.01, \*\*\**P*<0.001, and \*\*\*\**P*<0.0001).

## Author contributions

I.S. conceived the study. X.Z, Y.K., Y.L., Y.D., F.L., and M.M. performed various portions of the animal experiments, histologic analyses, and in vitro experiments. L.T. and B.W., performed EPA, DHA and resolvin metabolite profiling in colonic mucosa cells and sera by LC/MS/MS with guidance from P.Y. I.S. and X.Z. designed and guided the experiments, analyzed the data, and wrote the manuscript. D.W. and P.Y. provided conceptual feedback for the manuscript.

## Acknowledgments

This study made use of the MD Anderson Cancer Center Genetically Engineered Mouse Facility; the Research Animal Support Facility, supported by Cancer Center Support Grant P30CA016672; and the MD Anderson Cancer Center metabolomics facility service, supported in part by Cancer Prevention Research Institute of Texas (CPRIT) grant RP130397 and NIH grant 1S10OD012304-01. The manuscript was edited by Jennifer Peterson in the Department of Gastrointestinal Medical Oncology at The University of Texas MD Anderson Cancer Center.

## Ethical statement

This animal study was approved by the Institutional Animal Care and Use Committee of The University of Texas MD Anderson Cancer Center (IACUC #001434). All institutional and national guidelines for the care and use of laboratory animals were followed.

## Supplementary

**Supplementary Figure 1.**
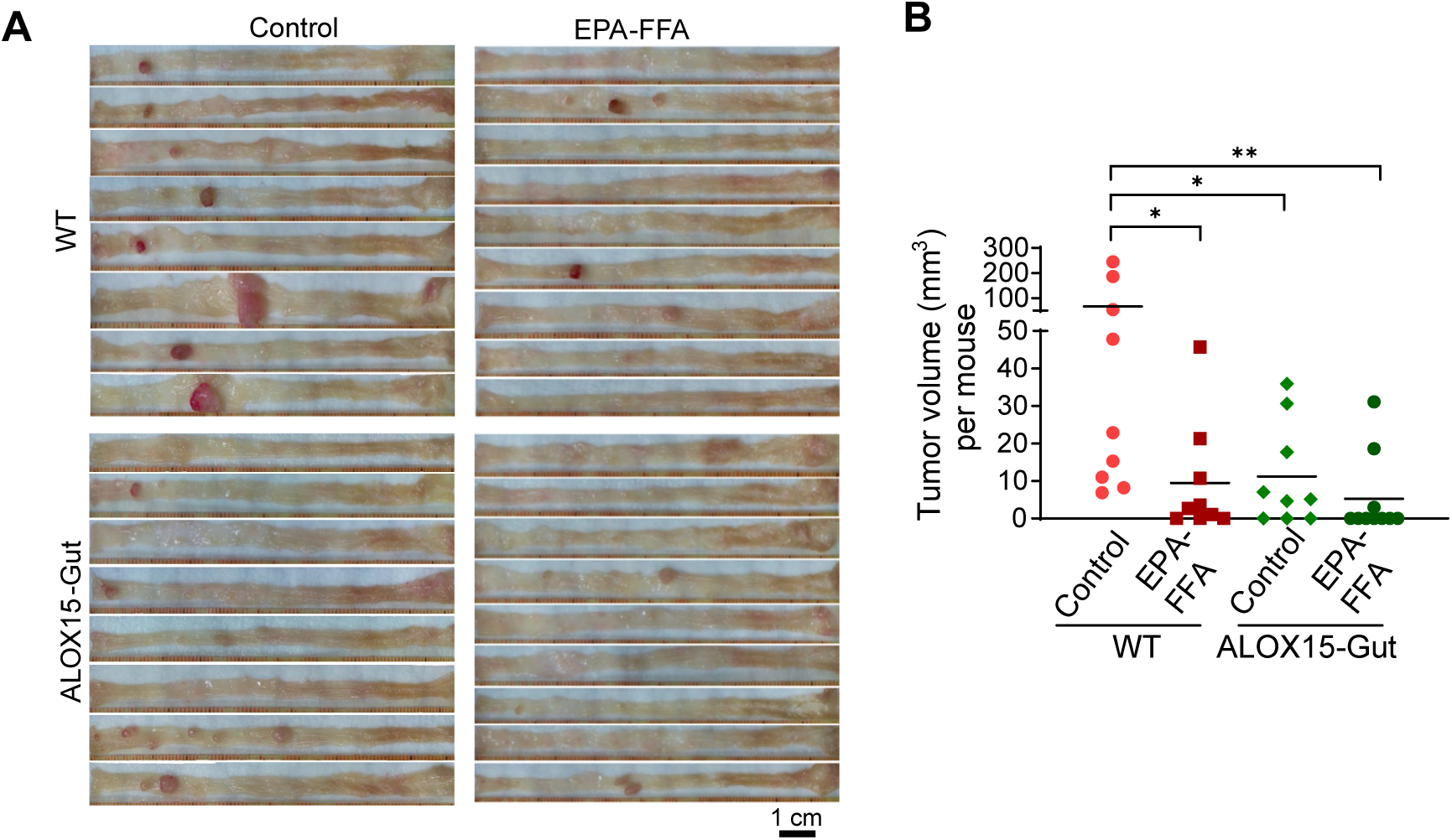
Effects of intestinal ALOX15 transgenic expression on EPA-FFA modulation of AOM-induced CRC. (A, B) ALOX15-Gut and their wild type (WT) littermates were fed either control diet or 1% EPA-FFA supplemented diet and treated with AOM (7.5 mg/kg once a week for 6 consecutive *i. p.* injections to induce CRC. The mice were euthanized at 20 weeks after the last AOM injection (n = 9-10 mice per group). (A) Representative gross colon pictures of the indicated mice. (B) Tumor volumes were measured. Lines for panel B represent means. Chi-square test for panel B. * *P < 0.05; ** P < 0.01*.

**Supplementary Figure 2.**
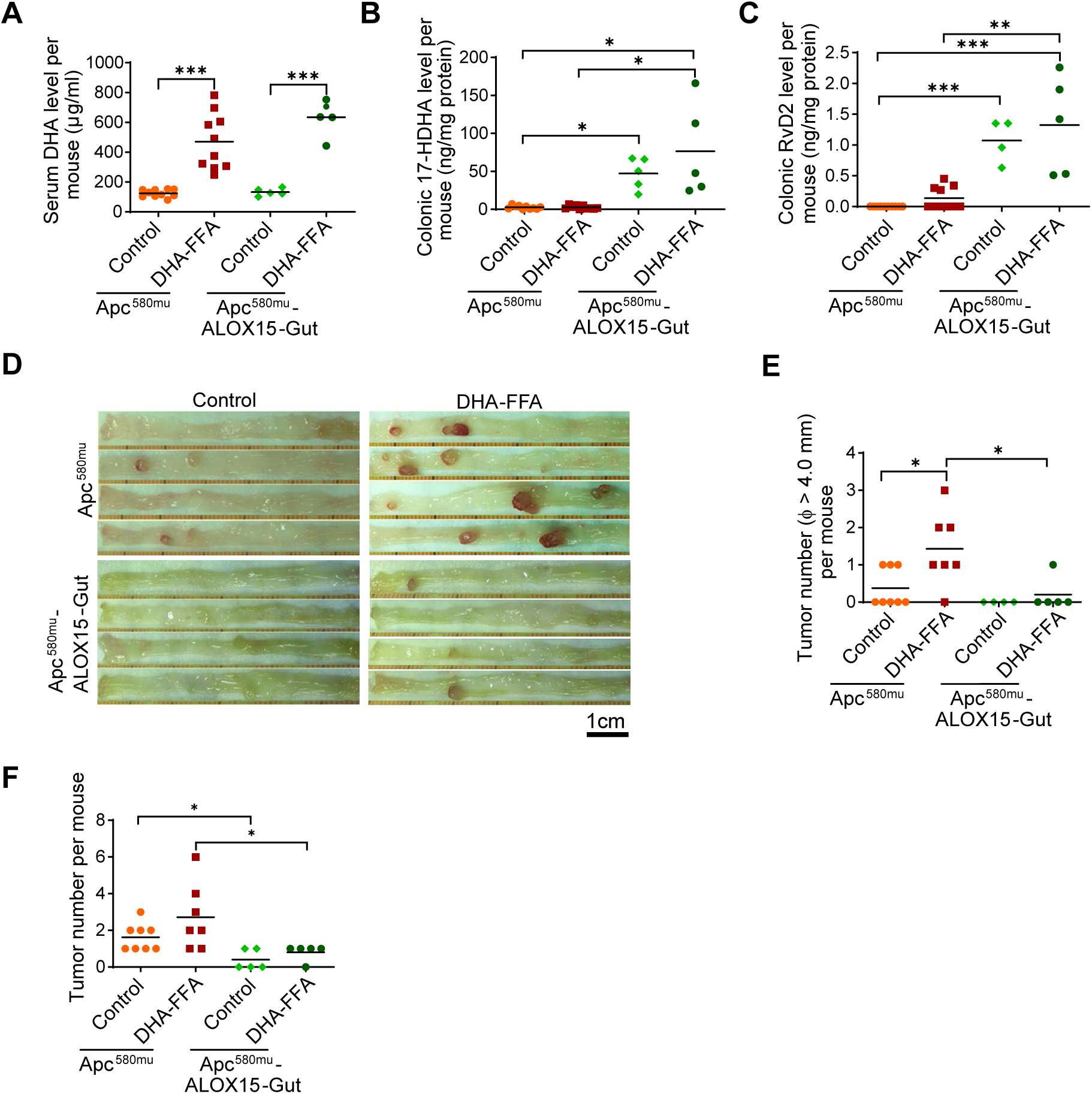
Effects of intestinal ALOX15 transgenic expression on DHA modulation of Apc mutation-induced CRC. Apc^580mu^-ALOX15-Gut and Apc^580mu^ mice at 4-6 weeks were fed either control diet or 1% DHA-FFA-supplemented diet for 10 weeks and then euthanized. Colonic tumor number per mouse was counted, and the blood and the scraped colonic mucosa cells were harvested (n = 6-8 mice per group). (A-C) Serum DHA (A), colonic 17-HDHA (B) and RvD2 (C) level per mouse for the indicated mouse groups, measured by LC/MS/MS. (D-F) Representative gross colon pictures (D), large colonic tumor number (diameter φ ≥ 4.0 mm) (E), and total colonic tumor number (F) per mouse for the indicated mouse groups. Lines for panels A-C and E-F represent means. Two-way ANOVA with Bonferroni adjustments for all multiple comparisons for panels A-C, and Chi-square test for panels E, F. *\* P < 0.05; *** P <0 .001*.

**Supplementary Figure 3.**
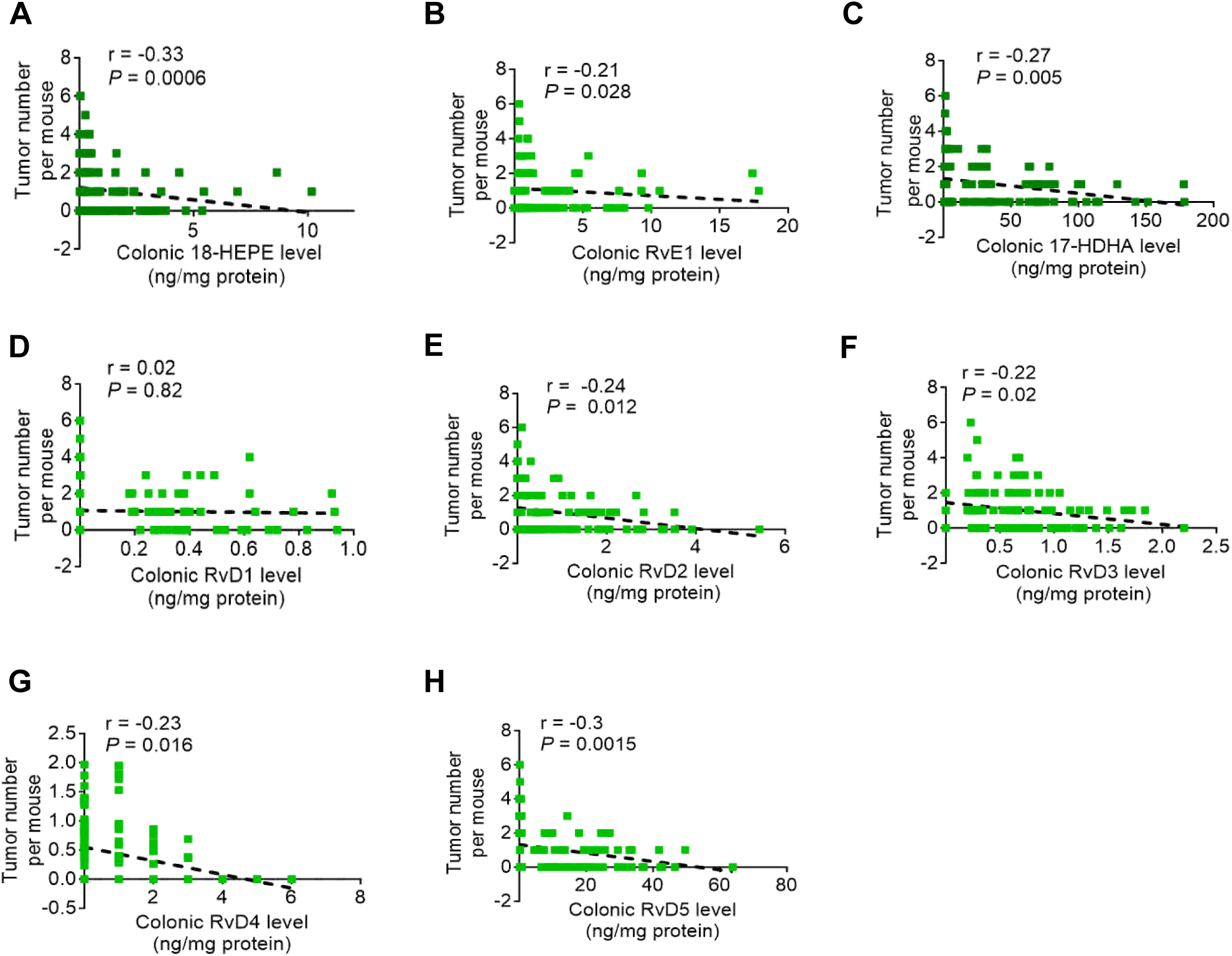
Correlation analysis of colonic tumor number with colonic levels of resolvins E and D series and their precursors. Spearman correlation analysis of colonic tumor number with colonic levels of 18-HEPE (A), RvE1 (B), 17-HDHA (C), and RvD1-5 (D-H) for total Apc^580mu^-ALOX15-Gut and Apc^580mu^ mice treated with EPA and DHA as described in Figure 3.

**Supplementary Figure 4.**
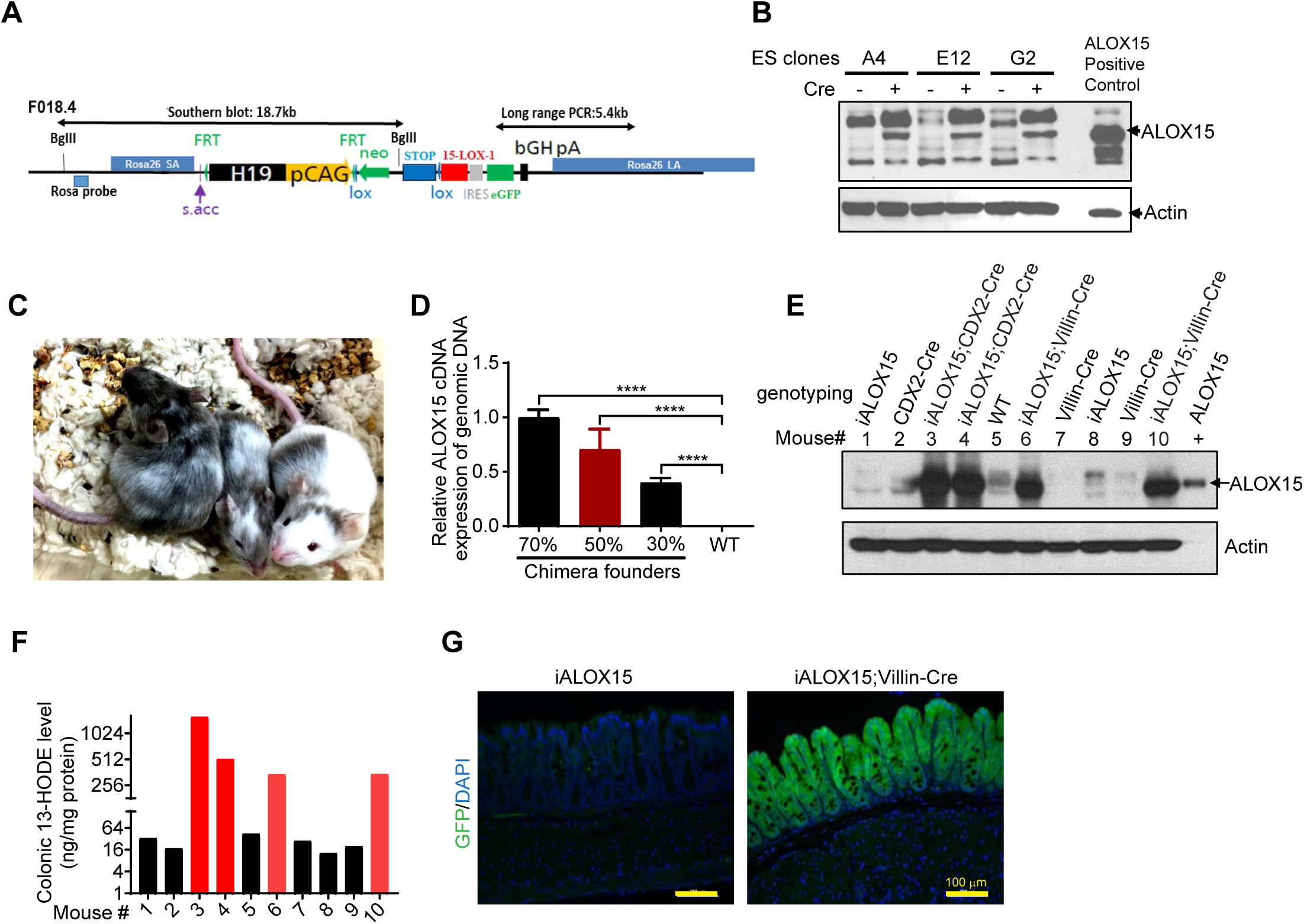
Generation and characterization of a conditional ALOX15 mouse model (iALOX15). (A) Schematic for the generation of iALOX15. Two mouse models could be generated from one construct as shown: the first allowing inducible ALOX15 expression driven by the CAG promoter, and the second was driven by the mouse Rosa26 promoter. The genuine Rosa26 promoter is further upstream, and once in embryonic stem (ES) cells, its transcript is spliced into the splice acceptor of exon 2 (“s.acc.”, purple). In the native state, transcription is aborted at the insulator site (H19, black), and instead starts at the CAG promoter (pCAG, yellow arrow). The loxP-flanked (lox, blue arrow heads) neo–STOP cassette for conditional activation has a stop site that is a large region of the SV40 intron plus late poly A (neo, green arrow; STOP, blue box). The ALOX15 cDNA is inserted downstream of the neo/stop cassette and is followed by an IRES-driven eGFP reporter gene. For exchange of the promoters, the CAG-cassette is flanked by FRT-sites. (B) ALOX15 expression of the three selected ES cell clones infected with or without Cre adenovirus. Three clones of ES cells (A4, E12 and G2) were cultured and infected with Cre adenovirus (MOI=200) for 24 hours. ALOX15 expression was measured by Western blot. (C, D) Generation and genotyping of iALOX15 chimeric mice. Two ES cell clones (A4, E12) were injected into blastocysts from C57BL/6 albino mice. Three of the living pups being 30-70% chimeric (black on white) were derived from ES cell clone A4 (C). ALOX15 cDNA expression level of genomic DNA was measured by qPCR. \*\*\*\**P < 0.0001* compared to C57BL/6 WT (WT: negative control) (D). (E-G) Characterization of the new iALOX15 mouse model. iALOX15 mice with Rosa26 promoter were bred with either CDX-Cre (mouse #1 to #4) or Villin-*Cre* recombinase mice (mouse #5 to #10) to induce ALOX15 expression via targeted intestinal Cre recombinase expression. (E) Colonic ALOX15 expression was observed in colonic samples from mice that carried both iALOX15 and Cre transgenic genes (mouse #3 and #4 [iALOX15;CDX-Cre], mouse #6 and #10 [iALOX15;Villin-Cre]) but not in those that just carried iALOX15 alone (mouse #1 and #8) or Cre gene alone (mouse #2 [CDX2-Cre], mouse #7 and #9 [Villin-*Cre*]) or none of them [wild-type (WT)] (mouse #5). + represents ALOX15 positive control. (F) Colonic ALOX15 expression was enzymatically active, as demonstrated by markedly increased 13-HODE. (G) Successful Cre recombinase in iALOX15; VIllin-Cre mice also induced eGFP expression, measured by immunofluorescence staining.

**Supplementary Figure 5.**
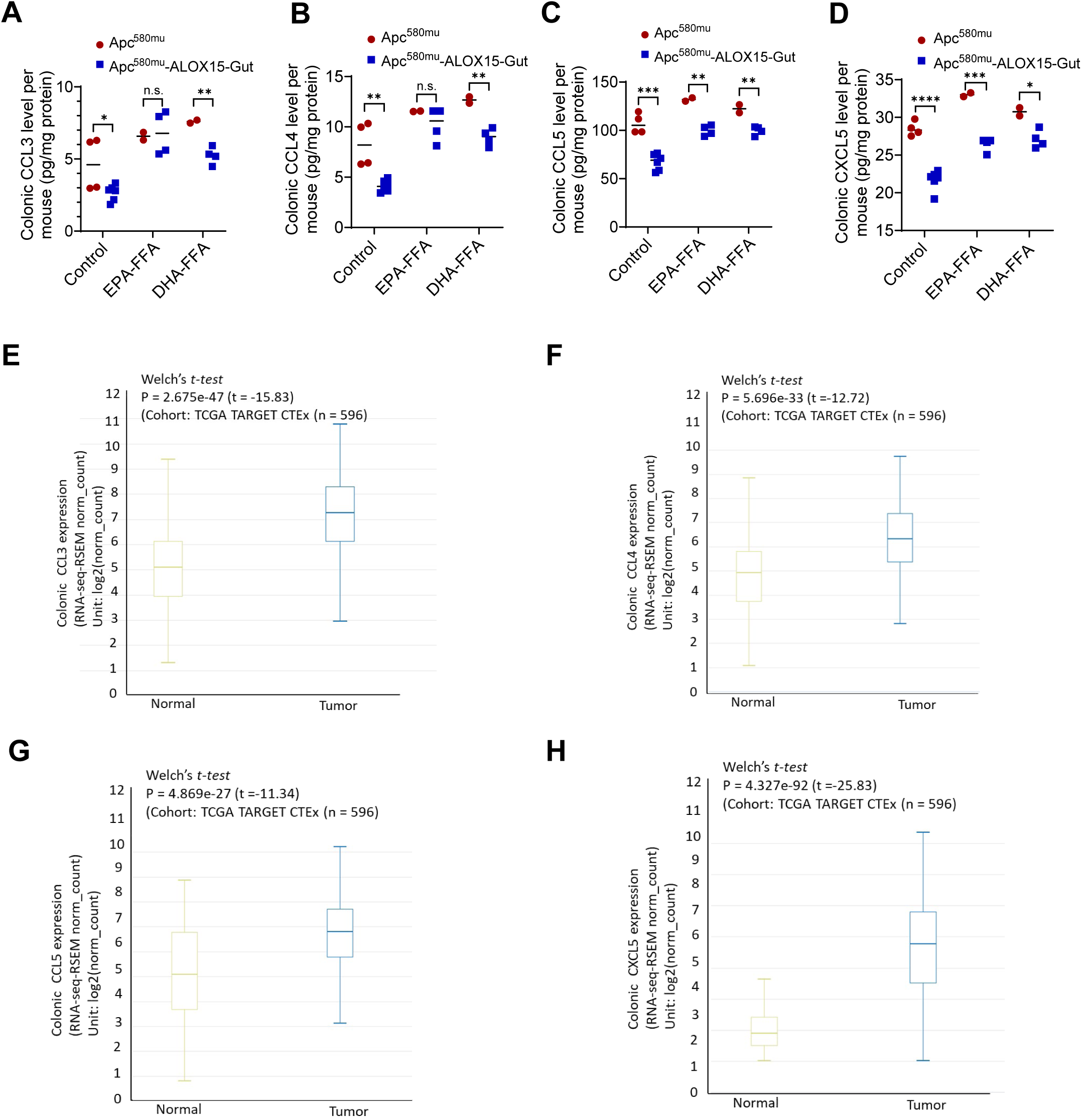
ALOX15 modulates the effects of EPA and DHA on the production of the pro-tumorigenic chemokines and cytokines. (A-D) Apc^580mu^-ALOX15-Gut and Apc^580mu^ mice at 6 weeks were fed 1% DHA-FAA diet, 1% EPA-FFA diet or control diet for 6 weeks and then euthanized. Quantitative results of CCL3 (A), CCL4 (B), CCL5 (C) and CXCL5 (D) protein levels in the scraped colorectal epithelial cells for the indicated mice. (E-H) Comparison of CCL3-5 and CXCL5 expression between human colorectal cancers and their adjacent normal colon tissues. USC Xena portal analyses of TCGA TARGET GTEx RNAseq data for CCL3 (E), CCL4 (F), CCL5 (G) and CXCL5 (H) mRNA expression levels in paired normal colon tissues and primary colorectal cancers (n=596).

**Supplementary Figure 6.**
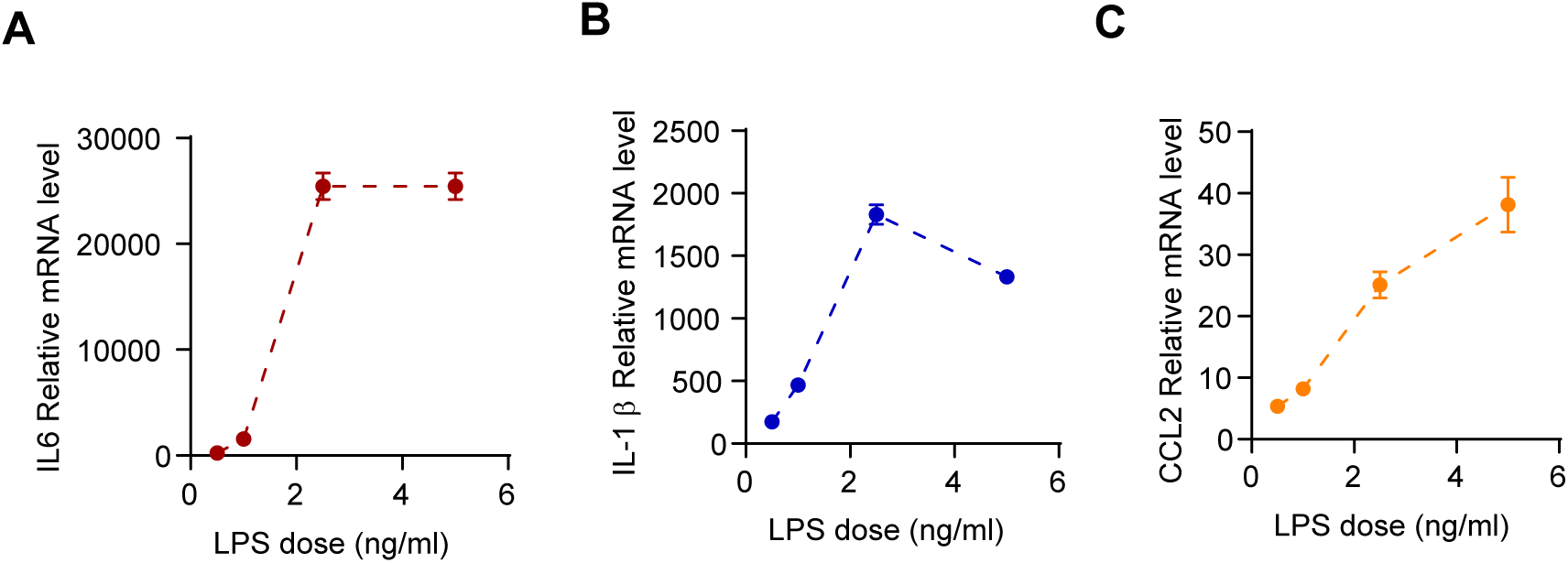
Effects of LPS on the expression of the cytokines in Raw 264.7 mouse macrophages. Raw 264.7 cells were treated with LPS at the indicated concentrations for 4 hours and then harvested for total RNA extraction. (A) IL-6, (B) IL-β and (C) CCL2 mRNA expression levels were measured by RT-qPCR. Data are mean ± SEM.

**Supplementary Table 1.** Formula for menhaden fish oil and control diets.

| **Product #** | **D12070201** |  | **D12070202** |  |
| --- | --- | --- | --- | --- |
| **Diet** | **control** |  | **Menhaden fish oil** |  |
|  | gm% | kcal% | gm% | kcal% |
| **Protein** | 20 | 19 | 20 | 19 |
| **Carbohydrate** | 56 | 50 | 56 | 50 |
| **Fat** | 15 | 31 | 15 | 31 |
| **Total** |  | **100** |  | **100** |
| **kcal/gm** | **4.4** |  | **4.4** |  |
| **Ingredient** | **gm** | **kcal** | **gm** | **kcal** |
| **Casein** | 200 | 800 | 200 | 800 |
| **DL-Methionine** | 3 | 12 | 3 | 12 |
| **Corn Starch** | 145 | 580 | 145 | 580 |
| **Maltodextrin 10** | 75 | 300 | 75 | 300 |
| **Sucrose** | 320 | 1280 | 320 | 1280 |
| **Cellulose, BW200** | 50 | 0 | 50 | 0 |
| **Corn Oil** | 150 | 1350 | 35 | 315 |
| **Menhaden Oil with 200 ppm tBHQ** | 0 | 0 | 115 | 1035 |
| **Mineral Mix S10001** | 35 | 0 | 35 | 0 |
| **Vitamin Mix V10001** | 10 | 40 | 10 | 40 |
| **Choline Bitartrate** | 2 | 0 | 2 | 0 |
| **BHT** | 0.2 | 0 | 0.177 | 0 |
| **tBHQ** | 0.2 | 0 | 0.2 | 0 |
| **FD&C Yellow Dye #5** | 0.05 | 0 | 0 | 0 |
| **FD&C Blue Dye #1** | 0 | 0 | 0.05 | 0 |
| **Total** | **990.45** | **4362** | **990.427** | **4362** |

**Supplementary Table 2.**
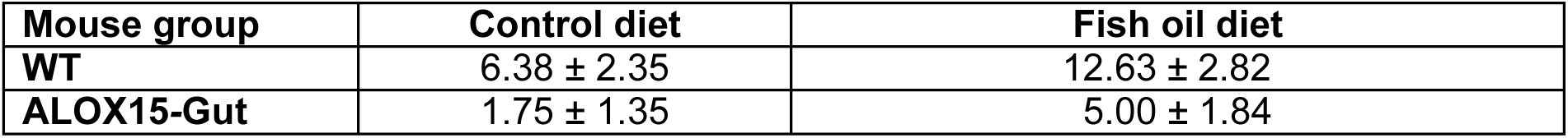
The effects of dietary menhaden fish oil and ALOX15 on DSS/AOM–induced colitis-associated colorectal cancer [tumor number per mouse (mean ± SEM)].

**Supplementary Table 3.** Formula for 1% EPA-FFA, 1% DHA-FFA, 2.7% Lovaza, 0.25% Lovaza and control diets.

| Product # | D03090904P |  | D12090801 |  | D12090806 |  | D12090803 |  | D20042001 |  |
| --- | --- | --- | --- | --- | --- | --- | --- | --- | --- | --- |
| Diet | Control |  | 1% EPA-FFA |  | 1% DHA-FFA |  | 2.7% Lovaza |  | 0.25% Lovaza |  |
|  | gm% | kcal% | gm% | kcal% | gm% | kcal% | gm% | kcal% | gm% | kcal% |
| Protein | 20 | 20 | 20 | 20 | 20 | 20 | 20 | 20 | 20 | 20 |
| Carbohydrate | 64 | 64 | 64 | 64 | 63 | 64 | 64 | 64 | 64 | 64 |
| Fat | 7 | 16 | 7 | 16 | 9 | 16 | 7 | 16 | 7 | 16 |
| Total |  | 100 |  | 100 |  | 100 |  | 100 |  | 100 |
| kcal/gm | 4 |  | 4 |  | 3.9 |  | 4 |  | 4 |  |
| Ingredient | gm | kcal | gm | Kcal | gm | kcal | gm | kcal | gm | kcal |
| Casein | 200 | 800 | 200 | 800 | 200 | 800 | 200 | 800 | 200 | 800 |
| L-Cystine | 3 | 12 | 3 | 12 | 3 | 12 | 3 | 12 | 3 | 12 |
| Corn Starch | 397.486 | 1589.9 | 397.486 | 1589.9 | 397.486 | 1589.9 | 397.486 | 1589.9 | 397.486 | 1589.9 |
| Maltodextrin 10 | 132 | 528 | 132 | 528 | 132 | 528 | 132 | 528 | 132 | 528 |
| Sucrose | 100 | 400 | 100 | 400 | 100 | 400 | 100 | 400 | 100 | 400 |
| Cellulose, BW200 | 50 | 0 | 50 | 0 | 50 | 0 | 50 | 0 | 50 | 0 |
| Soybean Oil | 0 | 0 | 0 | 0 | 0 | 0 | 0 | 0 | 0 | 0 |
| Corn Oil | 70 | 630 | 60 | 540 | 60 | 540 | 43 | 387 | 67.5 | 607.5 |
| EPA-FFA | 0 | 0 | 10 | 90 | 0 | 0 | 0 | 0 | 0 | 0 |
| DHA-FFA | 0 | 0 | 0 | 0 | 10 | 90 | 0 | 0 | 0 | 0 |
| Lovaza | 0 | 0 | 0 | 0 | 0 | 0 | 27 | 243 | 2.5 | 22.5 |
| tBHQ | 0.014 | 0 | 0.014 | 0 | 0.014 | 0 | 0.014 | 0 | 0.014 | 0 |
| Mineral Mix S10022G | 35 | 0 | 35 | 0 | 35 | 0 | 35 | 0 | 35 | 0 |
| Vitamin Mix V10037 | 10 | 40 | 10 | 40 | 10 | 40 | 10 | 40 | 10 | 40 |
| Choline Bitartrate | 2.5 | 0 | 2.5 | 0 | 2.5 | 0 | 2.5 | 0 | 2.5 | 0 |
| FD&C Yellow Dye #5 | 0 | 0 | 0 | 0 | 0 | 0 | 0.05 | 0 | 0.025 | 0 |
| FD&C Red Dye #40 | 0.025 | 0 | 0.05 | 0 | 0 | 0 | 0 | 0 | 0.025 | 0 |
| FD&C Blue Dye #1 | 0.025 | 0 | 0 | 0 | 0.05 | 0 | 0 | 0 | 0 | 0 |
| Total | 1000.05 | 4000 | 1000.05 | 4000 | 1000.05 | 4000 | 1000.05 | 4000 | 1000.05 | 4000 |

**Supplemental Table 4.** The effects of ALOX15 and dietary 1% EPA-FFA, 1% DHA-FFA, and 2.7% Lovaza on AOM–induced colorectal tumor volumes (mm^3^).

| Mouse group | Control diet | DHA-FFA diet | EPA-FFA diet | Lovaza diet |
| --- | --- | --- | --- | --- |
| WT | 15.75 ± 5.79 | 17.53 ± 5.06 | 7.16 ± 5.41 | 2.82 ± 1.46<sup>a</sup> |
| ALOX15-Gut<sup>+/-</sup> | 7.50 ± 2.72 | 7.17 ± 3.76 | 6.18 ± 3.70 | 0.19 ± 0.10 |
| ALOX15-Gut<sup>+/+</sup> | 3.70 ± 2.13<sup>b</sup> | 8.24 ± 3.34 | 5.80 ± 3.30 | 1.97 ± 0.81<sup>c</sup> |

- a. $P = 0.0036$ for the comparison of Lovaza diet–fed WT mice vs. control diet–fed WT mice. - b. $P = 0.019$ for the comparison of control diet–fed ALOX15-Gut<sup>+/+</sup> vs. control diet–fed WT mice. - c. $P = 0.0021$ for the comparison of Lovaza–fed ALOX15-Gut<sup>+/+</sup> vs. control diet–fed WT mice. - by two-way ANOVA with Bonferroni adjustments for all multiple comparisons.

**Supplementary Table 5.**
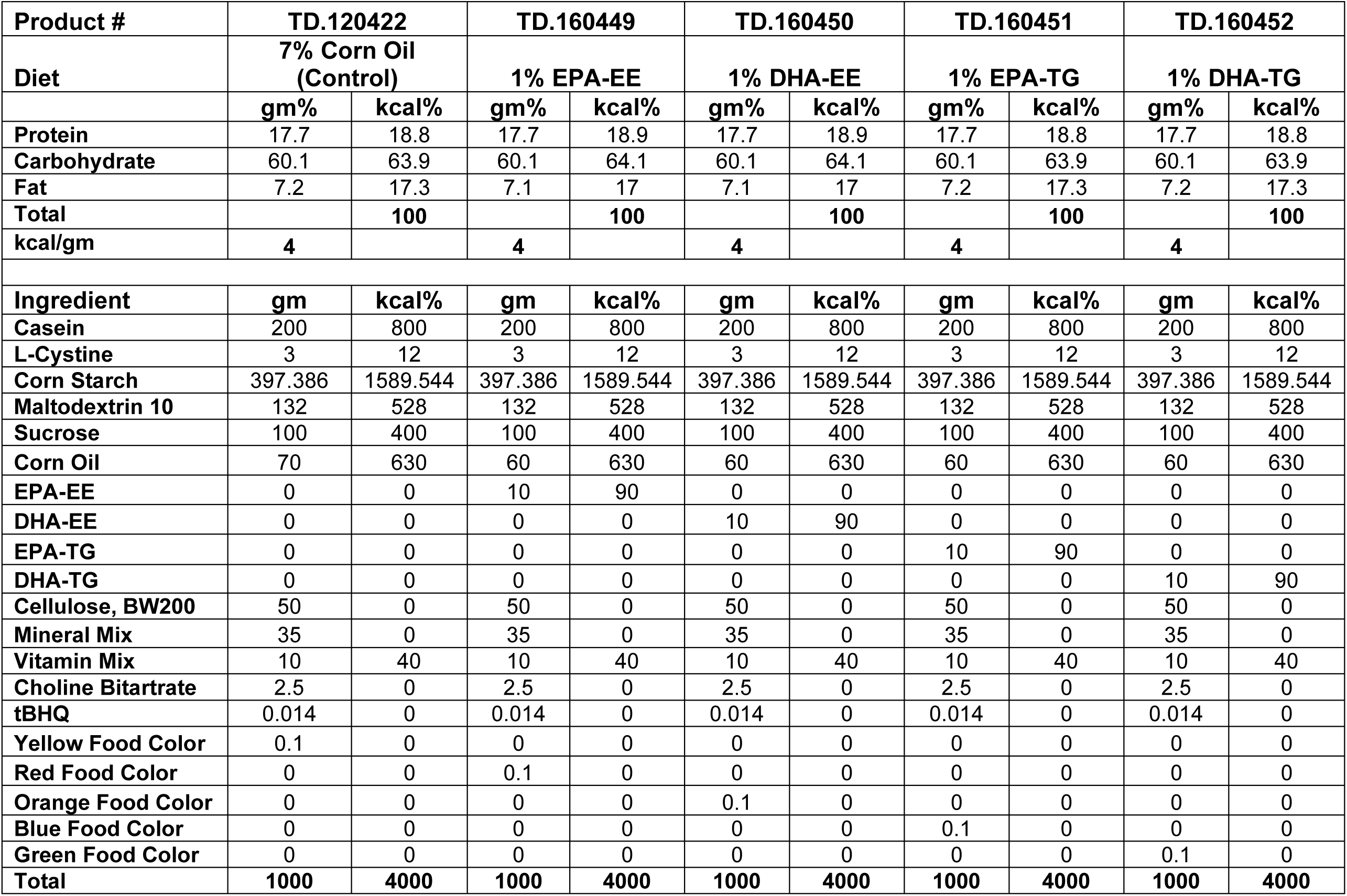
Formula for 1% EPA-EE, 1% EE-TG, 1% DHA-EE, 1% DHA-TG and control diets.

**Supplementary Table 6.** The effects of dietary 1% DHA and 1% EPA in EE and TG formulations on the production of colonic resolvins in Apc^580mu^ and Apc^580mu^-ALOX15-Gut mice.

| Diet | 18-HEPE (ng/mg protein) |  |  | RvE1 (ng/mg protein) |  |  | 17-HDHA (ng/mg protein) |  |  | RvD1 (ng/mg protein) |  |  |
| --- | --- | --- | --- | --- | --- | --- | --- | --- | --- | --- | --- | --- |
|  | *Apc*<sup>580mu</sup> | *Apc*<sup>580mu</sup>-ALOX15-Gut | *P* | *Apc*<sup>580mu</sup> | *Apc*<sup>580mu</sup>-ALOX15-Gut | *P* | *Apc*<sup>580mu</sup> | *Apc*<sup>580mu</sup>-ALOX15-Gut | *P* | *Apc*<sup>580mu</sup> | *Apc*<sup>580mu</sup>-ALOX15-Gut | *P* |
| Control | 0.03 | 0.04 | 0.987 | 0.36 | 0.48 | 0.872 | 1.33 | 22.74 | 0.0143 | 0.00 | 0.21 | 0.016 |
| DHA-EE | 0.26 | 1.94 | 0.004 | 0.75 | 2.36 | 0.0547 | 3.21 | 92.60 | <.0001 | 0.06 | 0.07 | 0.8697 |
| DHA-TG | 0.41 | 1.78 | 0.012 | 0.67 | 2.91 | 0.0046 | 4.26 | 101.41 | <.0001 | 0.56 | 0.28 | 0.0014 |
| EPA-EE | 0.96 | 4.41 | <.0001 | 0.42 | 9.24 | <.0001 | 1.49 | 55.30 | <.0001 | 0.26 | 0.43 | 0.0521 |
| EPA-TG | 0.39 | 3.02 | <.0001 | 0.48 | 5.68 | <.0001 | 1.53 | 39.18 | <.0001 | 0.00 | 0.10 | 0.2313 |
| Diet | RvD2 (ng/mg protein) |  |  | RvD3 (ng/mg protein) |  |  | RvD4 (ng/mg protein) |  |  | RvD5 (ng/mg protein) |  |  |
|  | *Apc*<sup>580mu</sup> | *Apc*<sup>580mu</sup>-ALOX15-Gut | *P* | *Apc*<sup>580mu</sup> | *Apc*<sup>580mu</sup>-ALOX15-Gut | *P* | *Apc*<sup>580mu</sup> | *Apc*<sup>580mu</sup>-ALOX15-Gut | *P* | *Apc*<sup>580mu</sup> | *Apc*<sup>580mu</sup>-ALOX15-Gut | *P* |
| Control | 0.05 | 0.59 | 0.0088 | 0.13 | 0.69 | <.0001 | 0.04 | 0.17 | 0.0333 | 0.04 | 10.05 | 0.0006 |
| DHA-EE | 0.07 | 1.75 | <.0001 | 0.54 | 1.20 | <.0001 | 0.00 | 1.19 | <.0001 | 0.45 | 30.96 | <.0001 |
| DHA-TG | 0.06 | 3.00 | <.0001 | 0.65 | 1.31 | <.0001 | 0.14 | 1.46 | <.0001 | 0.34 | 31.99 | <.0001 |
| EPA-EE | 0.00 | 1.39 | <.0001 | 0.31 | 0.79 | 0.0001 | 0.11 | 0.60 | <.0001 | 0.23 | 20.00 | <.0001 |
| EPA-TG | 0.07 | 0.62 | 0.0077 | 0.19 | 0.60 | 0.0007 | 0.00 | 0.47 | <.0001 | 0.12 | 12.26 | <.0001 |

\* *P* values are calculated by two-way ANOVA with Bonferroni adjustments for all multiple comparisons.

